# Muscle-derived Cues are Required to Specify Proprioceptor Pool Identity

**DOI:** 10.1101/2022.08.18.504408

**Authors:** Amy L. Norovich, Susan Brenner-Morton, Thomas M. Jessell

## Abstract

The formation of spinal sensory-motor circuits requires the diversification of proprioceptive sensory neurons (pSNs). During embryonic development, pSNs acquire molecular identities aligned with the limb muscle that they supply, but the extent of pSN “pool” diversity and how it is established are poorly understood. We find that the gene *v-set transmembrane domain-2b* (*vstm2b*) is preferentially expressed in pSN pools supplying dorsal limb muscle targets along the proximodistal extent of the limb. Genetic removal of muscle precursor cells from the developing limb greatly reduces the number of pSNs expressing *vstm2b*, demonstrating a requirement for limb muscle in specifying pSN pool identity. Comparison of dorsal and ventral muscle precursors identifies spatially restricted expression of the genes *lumican* (*lum*), *decorin* (*dcn*), and *BMP binding endothelial regulator* (*bmper*), demonstrating that dorsal and ventral muscle groups possess distinct molecular identities early in embryonic development. Together, these findings show that limb muscle is required for the specification of pSN pool identity and define early molecular correlates of dorsoventral muscle identity that are positioned to drive neuronal diversity.

## Introduction

The array of limb movements available to terrestrial animals is directed by sensory-motor circuits in the spinal cord. Each of the ∼60 muscles of the hindlimb is supplied by a dedicated pool of spinal motor neurons which, in turn, receive feedback about the state of that muscle from proprioceptive sensory neurons (pSNs) in the dorsal root ganglion (DRG). The ability of pSNs to refine motor neuron activity is rooted in the highly specific and conserved pattern of monosynaptic connections that they form with motor pools. During embryonic development, pSNs form strong connections with motor pools supplying the same limb muscle and weaker connections with those supplying muscles with similar biomechanical function, while avoiding contact with motor pools that supply muscles of opposing or unrelated function (Eccles et al., 1957; Frank and Westerfield, 1983; Hongo, 1984). This pattern of connectivity arises largely in the absence of patterned neural activity (Mears and Frank, 1997; Mendelsohn et al., 2015; Mendelson and Frank, 1991), implying that pSN pools possess distinct molecular identities that mediate selective synapse formation.

While the specification of motor neuron subtype identity is understood in considerable detail, less is known about how their sensory counterparts are organized and diversified (Imai and Yoshida, 2018; Stifani, 2014). Whereas motor neurons acquire subtype identities prior to the innervation of limb muscle targets, pSNs project into the limb with only generic sensory class specified (Imai and Yoshida, 2018). Recent studies have uncovered molecular signatures of pSN “pool” identities analagous those exhibited by motor neurons. Combinatorial expression of the genes *cdh13, sema5a*, and *crtac1* defines pSN pools occupying defined positions along the dorsoventral and proximodistal limb axes (Poliak et al., 2016), revealing an underlying positional logic to the specification of pSN pool identity. However, expression of these genes is restricted to pSNs supplying distal limb musculature, indicating that our characterization of pSN pool diversity is incomplete. Moreover, expression of these genes begins following limb innervation (Poliak et al., 2016), suggesting that extrinsic limb signaling mechanisms are responsible for imparting muscle-specific pool identities in pSNs.

Surgical manipulation of chick limb bud has demonstrated the involvement of peripherally-derived cues in patterning central sensory-motor connections (Wenner and Frank, 1995). In line with this view, mouse genetic manipulations identified limb mesenchyme as a source of cues driving pool-specific patterns of gene expression in pSNs, which in turn influence specificity within individual sensory-motor reflex arcs (Poliak et al., 2016). It is unclear, however, whether limb mesenchyme alone contains sufficient patterning information to instruct the full array of pSN subtype diversity required to achieve selective connectivity with the ∼60 motor pools that supply muscles of the hindlimb. To this end, muscle itself has been proposed to act as an additional source of pSN patterning information (Wenner and Frank, 1995). Indeed, aspects of pSN differentiation and subtype specification depend on cues from skeletal muscle. During embryonic development, muscle spindle formation by type Ia and II proprioceptive afferents requires the interaction of pSN neuregulin1 and its receptor Erbb2 on intrafusal muscle fibers (Leu et al., 2003). At postnatal stages, muscle-by-muscle variation in the level of NT3 signaling through the sensory TrkC receptor drives distinct molecular responses in pSN subsets (de Nooij et al., 2013). However, molecular evidence that muscle instructs pSN pool identity is lacking. Moreover, whether muscles differ in molecular composition sufficiently early in embryonic development to influence pSN pool diversification is unknown.

To address these issues, we set out to identify additional genes that define pSN pools. Our screen of pSNs supplying the tibialis anterior (TA) and gastrocnemius (GS) muscles of the mouse hindlimb (Poliak et al., 2016) – shank muscles that exert opposing forces on the ankle – predicted differential expression of the gene *v-set transmembrane domain-2b* (*vstm2b*), a transmembrane immunoglobulin (Ig) superfamily protein. Here, we characterize expression of *vstm2b* in pSN pools and find that it preferentially marks pSNs supplying dorsal muscles along the proximodistal extent of the limb. Moreover, in contrast to previously identified pSN pool markers *cdh13, sema5a*, and *crtac1*, we find that *vstm2b* expression among DRG sensory populations is exclusive to pSNs. Genetic manipulation of individual tissue types during mouse limb development shows that pSN *vstm2b* expression requires the presence of muscle in the developing limb, revealing a novel requirement for muscle in patterning neuronal subtype identity.

We reasoned that if muscle acts to instruct pSN pool identity, embryonic muscle cells would exhibit molecular distinctions aligned with the pattern of pSN *vstm2b* expression. We performed a molecular screen comparing dorsal and ventral muscle precursors at the level of the shank, where *vstm2b* is expressed exclusively by dorsally-innervating pSNs, and found that the expression of three genes – *lum, dcn*, and *bmper* – was restricted to dorsal or ventral muscles around the time pSN axons invade the hindlimb. Expression of these genes matches the pattern of pSN *vstm2b* expression, suggesting that these genes may act to pattern pSN pool diversity. Taken together, our data demonstrate a role for muscle in specifying pSN pool identity and reveal early molecular distinctions between dorsal and ventral muscles that are positioned to drive neuronal gene expression.

## Results

### *vstm2b* expression defines muscle-type proprioceptors

To define molecular distinctions between pSNs supplying dorsal and ventral muscles acting at the same joint, we performed an RNA-Sequencing (RNA-Seq)-based screen comparing gene expression in pSNs supplying the tibialis anterior (TA; dorsal) and gastrocnemius (GS; ventral) muscles of the mouse hindlimb (Poliak et al., 2016; see Figure S1). We identified DRG neurons as likely pSNs by their expression of parvalbumin (Pv), a generic marker of all pSNs that also labels a small subset of low threshold cutaneous mechanoreceptors, in *Pv*::YFP mice (de Nooij et al., 2013; Poliak et al., 2016). We labeled pSN cell bodies according to limb muscle target via injection of the retrograde tracer cholera toxin B subunit/Alexa Fluor 555 (ctb^555^) into TA or GS muscles of *Pv*::YFP mice at postnatal day 0 (P0). We then isolated individual YFP^+^, ctb^555+^ pSNs from dissociated lumbar DRG and pooled them to generate 30-cell TA and GS pSN samples for RNA-seq and differential gene expression analysis (Poliak et al., 2016).

To further probe the nature of pSN pool identity, we turned to this differential expression analysis to identify additional candidate genes predicted to have an all-or-none difference between TA and GS pSNs. We focused on cell surface molecules due to their potential to mediate the formation of selective connections between pSNs and spinal motor pools. We identified *vstm2b*, a transmembrane immunoglobulin (Ig)-domain protein, as expressed highly in TA pSNs but at insignificant levels in GS pSNs (fold change = 24, p = 2.33×10^−10^; Figure 1A). In situ hybridization histochemistry (ISH) in lumbar DRG at P1 revealed high-level expression of *vstm2b* in a fraction of large-diameter neuronal cell bodies (Figure 1B), consistent with expression in a subset of hindlimb-innervating pSNs.

**Figure 1.**
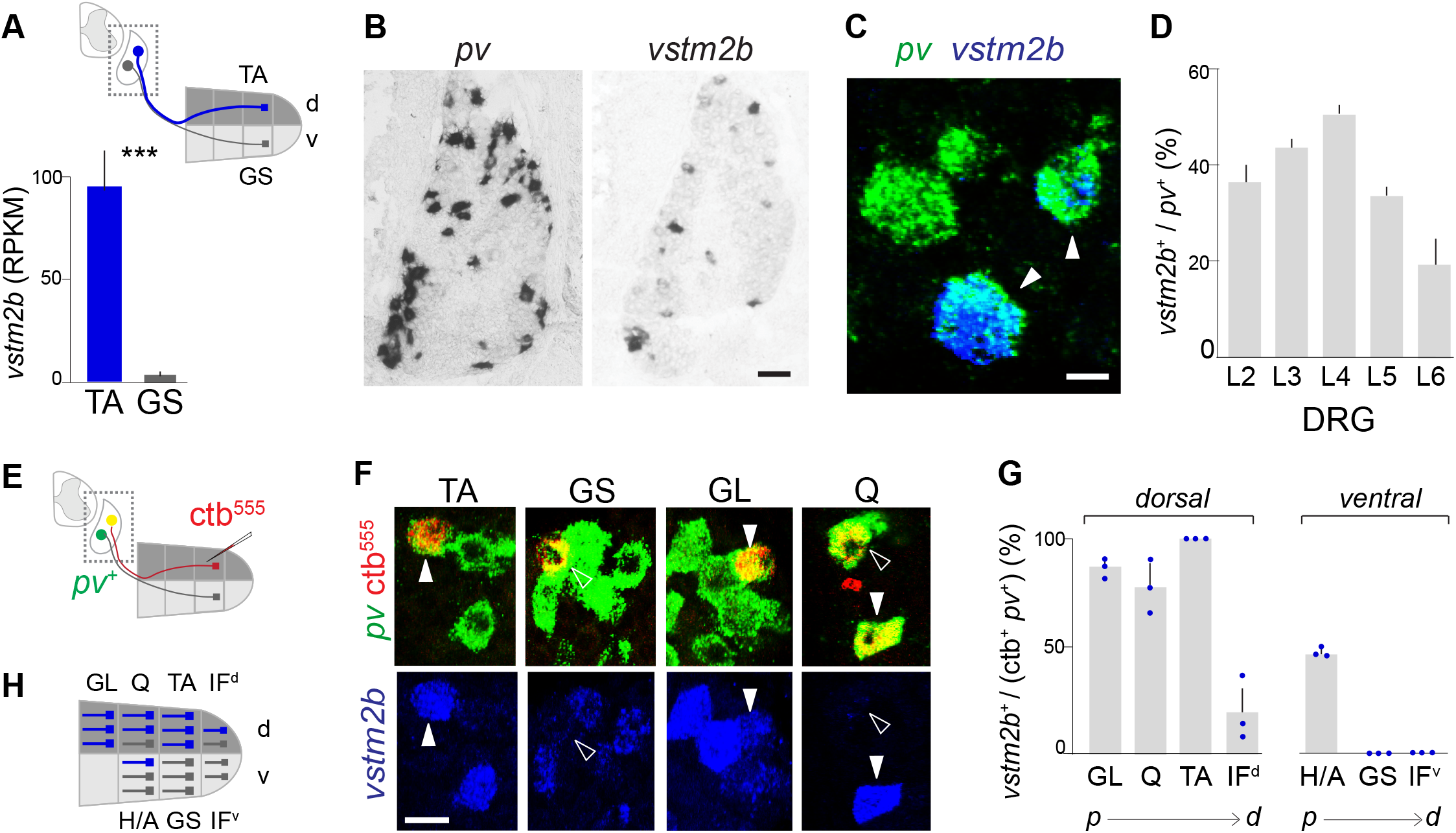
Expression of *vstm2b* in DRG neurons. (A) V-set transmembrane domain 2b (*vstm2b*) RNA expression in proprioceptive sensory neurons (pSNs) supplying the tibialis anterior (TA; d, dorsal) and gastrocnemius (GS; v, ventral) muscles of the shank at postnatal day 1 (P1) measured by RNA-Sequencing. Expression level is indicated in reads per kilobase per million (RPKM). Error bars represent SD from three biological replicates. ***, significant difference between TA and GS samples (p < 10^−9^). See also Figure S1. (B) In situ hybridization (ISH) showing parvalbumin (*pv*) expression by pSNs and *vstm2b* expression in a subset of large-diameter dorsal root ganglion (DRG) neurons. DRG L4 (L, lumbar) is shown. Scale bar, 40µm. (C-D) Subset of *pv*^+^ DRG neurons express *vstm2b* at P1. (C) Fluorescence *in situ* hybridization (FISH) for *vstm2b* and *pv*. Scale bar, 10µm. (D) Fraction of *pv*^+^ neurons expressing *vstm2b* in each lumbar DRG. Error bars represent SD; n=4, >200 neurons/level. (E-G) *vstm2b* status of muscle-type pSN subsets defined by peripheral termination. (E) Muscle-type pSNs were identified by ctb backfill of a muscle or muscle group, followed by FISH for *pv*. (F) *vstm2b* expression status of pSNs supplying the muscles or muscle groups tibialis anterior (TA), gastrocnemius (GS), gluteus group (GL), and quadriceps group (Q) at P1. Arrowheads indicate neurons defined as muscle-type pSNs by *pv* expression and ctb backfill of the indicated muscle. Closed: *vstm2b*^+^ pSNs; open: *vstm2b*^-^ pSNs. Scale bar, 20µm. (G) Fraction of GL, Q, TA, dorsal intrinsic foot (IF^d^), hamstring/adductor group (H/A), GS, and ventral intrinsic foot (IF^v^) pSNs expressing *vstm2b* (all groups: n=3). Data are ordered according to dorsoventral (d, v) and proximodistal (p=proximal; d=distal) position of each muscle or muscle group within the limb; represented as mean ± SD. (H) Schematic summarizing the *vstm2b* status of pSNs supplying muscles/muscle groups along the dorsoventral and proximodistal axes of the hindlimb.

Previously identified pSN pool markers *cdh13, sema5a*, and *crtac1* are expressed in several *pv*^-^ DRG sensory lineages in addition to *pv*^+^ pSNs (Poliak et al., 2016). We therefore sought to determine which populations of DRG sensory neurons express *vstm2b*, by examining the coincidence of *vstm2b* and *pv* via double fluorescence in situ hybridization (FISH) at P1. We found that 42% of L2-L6 *pv*^+^ DRG neurons expressed *vstm2b*, with the rostrocaudal distribution of *vstm2b*^*+*^ neurons increasing from 37% in L2 to a maximum of 51% in L4, then declining caudally to 20% in L6 (Figures 1C and 1D). Moreover, in contrast to other pSN genes, we observed that *vstm2b* labeling was confined to *pv*^+^ DRG neurons (Figures 1C and 1F), consistent with restriction of *vstm2b* expression to pSNs.

Seeking to confirm the all-or-none expression difference predicted by our RNA-Seq data, we next examined *vstm2b* expression in pSNs supplying TA and GS muscles. We injected ctb^555^ into TA or GS muscle at P0 and used FISH to evaluate *vstm2b* expression in *pv*^+^ ctb^555^-labeled pSNs at P1 (Figure 1E). We found that 100% of TA pSNs expressed *vstm2b*, whereas GS pSNs were devoid of *vstm2b* transcript (Figures 1F and 1G), confirming that *vstm2b* expression selectively marks TA but not GS pSNs.

TA pSN cell bodies are distributed among DRG L3-L5 (unpublished observation). Together, these three ganglia contain ∼520 *vstm2b*^+^ *pv*^+^ neurons – far more than the ∼60 pSNs estimated to supply TA muscle (Hunt, 1974; Poliak et al., 2016). *Vstm2b* must therefore be expressed in additional pSNs supplying other hindlimb muscles. To identify which muscles are supplied by *vstm2b*^+^ pSNs, we retrogradely labeled pSN cell bodies by injecting ctb^555^ into hip, thigh and foot muscles located in distinct dorsoventral and proximodistal domains of the hindlimb. Whereas TA and GS muscles are large and anatomically accessible, muscles of the hip, thigh and foot are nearly impossible to inject without spillover of ctb into adjacent muscles. We therefore chose to inject muscle synergy groups rather than individual muscles at these joints. For the proximodorsal gluteal group (GL; gluteus maximus and gluteus medius), we observed that nearly all (89%) backfilled *pv*^+^ pSNs expressed *vstm2b* (Figures 1F and 1G). At the thigh, we found that 72% of pSNs supplying the dorsal quadriceps group (Q; rectus femoris, vastus medius, vastus lateralis, and vastus intermedius) expressed *vstm2b*, and 42% of pSNs supplying the ventral hamstring/adductor group (H/A; gracilis, semitendinosus, semimembranosus, and adductor muscles) were marked by *vstm2b* transcript (Figures 1F and 1G). Most distally, the anatomy of intrinsic foot (IF) muscles precluded strict separation of dorsal from ventral musculature. Nevertheless, dorsally directed ctb^555^ injections revealed *vstm2b* expression in ∼25% of retrogradely labeled IF^d^ pSNs, whereas 0% of IF^v^ pSNs labeled by ventrally directed injections expressed *vstm2b* (Figure 1F).

Together, these findings indicate that dorsal muscles at all proximodistal points of the hindlimb are supplied preferentially by *vstm2b*-expressing pSNs (Figure 1H). Thus, there is a clear link between pSN *vstm2b* expression and dorsoventral termination within the hindlimb.

### Mapping proprioceptor Vstm2b status at peripheral terminals

The mixed *vstm2b* status observed for several muscle groups via ISH and backfill raised the issue of whether muscles within a synergy group have uniform but distinct *vstm2b* pool identities, or whether individual pSN pools are themselves mosaic in their *vstm2b* expression. We therefore sought to examine the status of individual pSN pools supplying each muscle within the groups profiled. To obtain a more detailed view of the *vstm2b* status of individual pSN pools, we generated an antibody to characterize Vstm2b protein localization in cell bodies and at peripheral terminals.

We first characterized the pattern of Vstm2b immunolabeling in lumbar DRG cell bodies. We assessed Vstm2b immunoreactivity in P1-3 mice with respect to immunolabeling for Pv and the transcription factor Rx3, the coincidence of which defines the pSN lineage (de Nooij et al., 2013). In each lumbar ganglion, Vstm2b labeling was observed in a subset of Pv^+^ Rx3^+^ DRG neurons (Figures 2A and 2B). Moreover, Vstm2b immunolabeling was restricted to Pv^+^ Rx3^+^ neurons (Figures 2A and 2B; Figure S2A), confirming that among DRG neurons, Vstm2b is restricted to pSNs. The fraction of Vstm2b^+^ pSNs observed in DRG L2-L6 matched the distribution observed by ISH, rising from 25% at L2 to a peak of 49% at L4, then declining caudally to 16% at L6 (Figure 2B). Thus, the distribution of lumbar DRG neurons expressing Vstm2b protein aligns with that observed for *vstm2b* transcript.

**Figure 2.**
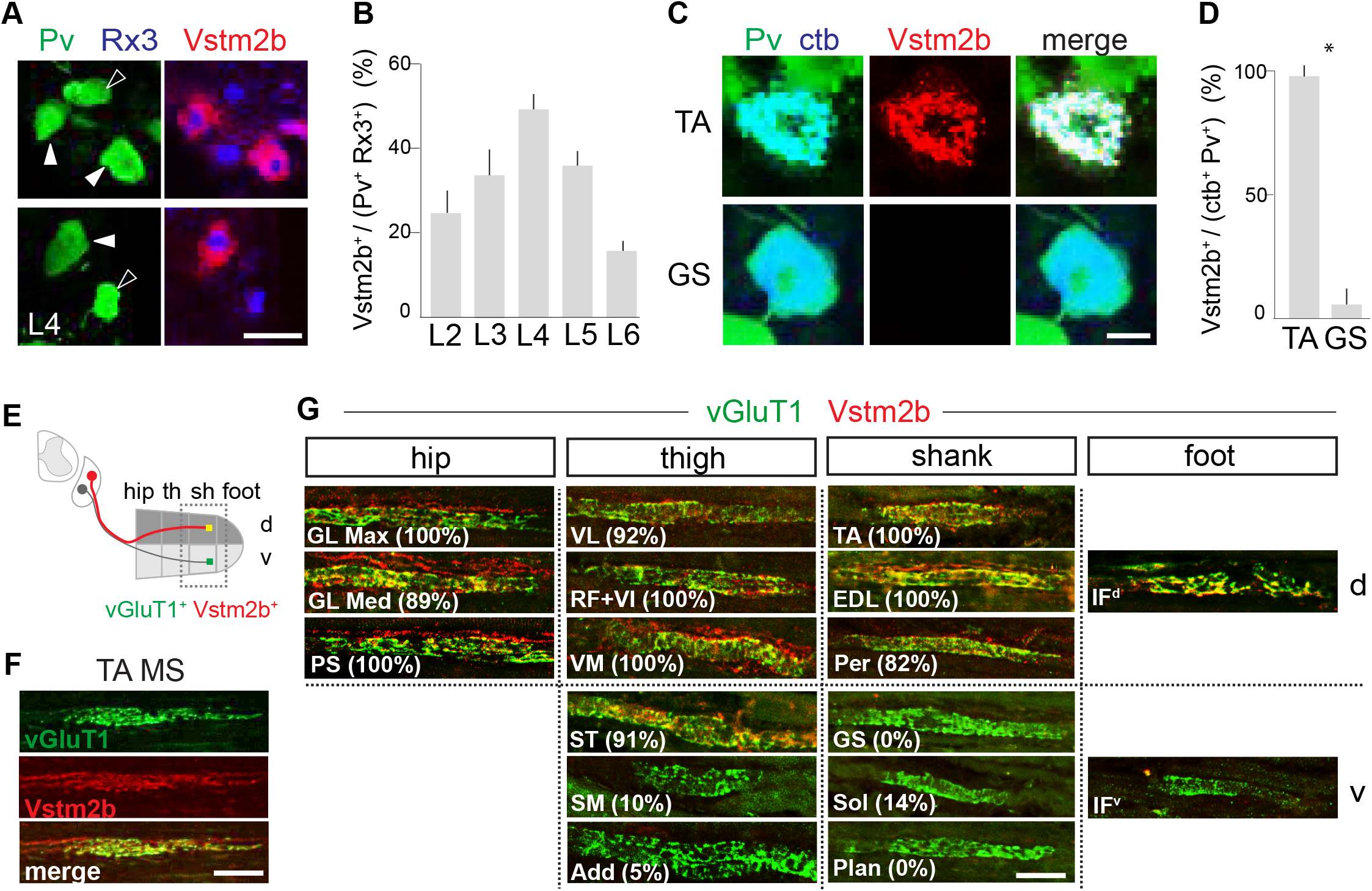
Mapping Vstm2b Protein Expression in pSNs Supplying Individual Hindlimb Muscles. (A-B) Vstm2b antibody labeling is restricted to Pv^+^ Rx3^+^ pSNs. (A) Immunohistochemistry in P3 L4 DRG. Arrowheads indicate pSNs, as defined by the coincidence of Pv and Rx3 immunolabeling. Closed: Vstm2b^+^ pSNs; open: Vstm2b^-^ pSNs. Scale bar, 30µm. (B) Fraction of Vstm2b^+^ pSNs in lumbar DRG at P1-3 (n=3; >100 neurons/level). Error bars represent SD. See also Figure S2A. (C-D) Vstm2b immunolabeling of muscle-type pSN cell bodies. (C) Representative images of Vstm2b immunolabeling of TA and GS pSNs identified by ctb backfill and Pv labeling at P3. Scale bar, 10µm. (D) Fraction of Vstm2b^+^ TA and GS group pSNs (TA: n=4 mice, GS: n=5 mice). Error bars represent SD. *, significant difference between TA and GS samples (Student’s t test, p < 10^−^ 6). (E-G) Vstm2b immunolabeling of muscle spindle sensory endings. (E) Vstm2b status of sensory endings was assessed by dissecting out individual muscles along the proximodistal and dorsoventral (d, v) limb axes and immunostaining for vGluT1 and Vstm2b. th, thigh; sh, shank. (F) Vstm2b labels TA muscle spindle (MS) terminals. P5 TA muscles immunostained for vGluT1 and Vstm2b. Scale bar, 50µm. (G) Vstm2b^+^ proprioceptors preferentially supply dorsal hindlimb muscles. Hindlimb muscles along the dorsoventral (d, v) and proximodistal limb axes were assessed for Vstm2b immunolabeling at spindle terminals in P3-5 mice (n ≥ 3 mice/muscle). Fraction of vGluT1^+^ muscle spindles colabeled by Vstm2b antibody is noted in parenthesis for each muscle. Scale bar, 50µm. See also Figures S2B-D.

We next examined the Vstm2b protein in defined pSN pools accessible by ctb backfill. *Vstm2b* transcript is observed in all TA and no GS pSNs (Figure 1), and we expected this pattern to be maintained by Vstm2b antibody labeling. At P1, we found that 98% of ctb^555^-labeled TA pSNs were marked by Vstm2b. In contrast, only 4% of GS pSNs were Vstm2b^+^, with rare Vstm2b-labeled pSNs presumed due to slight ctb spillover into adjacent muscles. Thus, we conclude that Vstm2b protein expression maintains the muscle-type distinction in *vstm2b* transcript expression observed for TA and GS pSNs and is faithfully represented in cell bodies by our Vstm2b antibody.

We proceeded to examine Vstm2b protein localization to peripheral pSN terminals. If present at sensory endings, examination of Vstm2b protein would allow us to characterize expression in pSN pools on a muscle-by-muscle basis as well as expression by individual neurons within each pool. Detection of *vstm2b* transcript and protein in 100% of identified TA pSN cell bodies (Figures 1G and 2D) suggests that all three functional subtypes of pSNs – groups Ia, II and Ib afferents – express *vstm2b*. To explore this, we examined Vstm2b immunolabeling at peripheral pSN terminals defined by vGluT1 antibody, which labels all pSN terminals in muscle (Figure 2E). In TA muscle, vGluT1^+^ terminals with the annulospiral morphology characteristic of group Ia and II muscle spindle (MS) afferents were marked by Vstm2b immunolabeling, demonstrating that Vstm2b protein localizes to peripheral terminals. We also noted Vstm2b^+^ vGluT1^+^ terminals with the flower spray morphology typical of type Ib Golgi tendon organ (GTO) afferents (Figure S2B), indicating that all pSN subtypes terminating in TA muscle share a common Vstm2b expression status. In contrast, all vGluT1^+^ sensory terminals within GS muscle lacked Vstm2b labeling (Figures 2G and S2B). Combined, these results show that Vstm2b protein localizes to peripheral terminals and preserves the Vstm2b status observed in muscle-type pSN cell bodies, with MSs and GTOs sharing a common status in TA and GS muscle.

We next extended this analysis to all muscles of the mouse hindlimb. We examined Vstm2b immunolabeling of MSs in individual muscles spanning the dorsoventral and proximodistal limb axes in P5 mice (Figure 2G). For muscles in dorsal shank, where TA muscle is located, we found that 100% of TA and extensor digitorum longus (EDL) sensory endings as well as 82% of peroneal group (PER: peroneus longus, brevis, and tertius) endings were marked by Vstm2b (Figures 2G and S2C). In contrast, ventral shank muscles GS, plantaris (Plan), flexor hallucis longus (FHL), and tibialis posterior (TP) were devoid of Vstm2b^+^ endings, and terminals in soleus (Sol) and flexor digitorius longus (FDL) muscles largely lacked Vstm2b labeling (Sol: 14% Vstm2b^+^, FDL: 17% Vstm2b^+^; Figures 2F and S1B). These results are again in agreement with our ISH experiments, where we observed that 100% of TA pool and 0% of GS pool pSNs expressed *vstm2b* (Figures 1G and S2D).

We proceeded to characterize Vstm2b status at muscle spindles in individual hip, thigh, and foot muscles. At the most proximal level of the hip, the vast majority of sensory endings found in dorsal muscles were marked by Vstm2b: 100% in gluteus maximus (GL Max), 89% in gluteus medius (GL Med), and 100% in psoas (PS; Figures 2G and S1C). Among dorsal thigh muscles, 92% of vastus lateralis (VL) and 100% of rectus femoris, vastus intermedius, and vastus medialis (RF, VI, and VM) sensory terminals had Vstm2b labeling (Figures 2G and S1C). Among ventral thigh muscles, only 10% semimembranosus (SM) and 5% adductor group (Add: adductors longus, brevis and magnus) sensory endings were Vstm2b^+^. However, large cohorts in ventral thigh muscles semitendinosus, gracilis, and biceps femoris were Vstm2b^+^ (ST, 91%; Grac, 100%; BF, 83%; Figures 2G and S1C). Finally, although we were unable to dissect dorsal from ventral intrinsic foot (IF) muscles prior to immunolabeling, we noted that 21% of all IF sensory endings received Vstm2b^+^ afferent innervation, where all Vstm2b^+^ afferents terminated in dorsal foot.

This muscle-by-muscle analysis shows that the vast majority of Vstm2b^+^ sensory endings are found in dorsal muscles, consolidating the view that Vstm2b^+^ pSNs preferentially supply dorsally-derived muscles along the extent of the hindlimb proximodistal axis.

### Proprioceptor *vstm2b* expression begins after limb innervation

The finding that expression of pSN pool markers *cdh13, sema5a*, and *crtac1* is assigned by limb mesenchyme (Poliak et al., 2016) led us to examine whether *vstm2b* expression is also induced by a peripheral cue. If pSN *vstm2b* expression is induced by a peripheral signal, then *vstm2b* transcript would be detectable only after sensory axons have entered the limb bud.

We examined the onset of *vstm2b* expression in lumbar pSNs relative to the timing of hindlimb innervation. Sensory nerves arrive at the lumbar plexus at e10.5 and establish dorsal and ventral branches within limb mesenchyme by e11.5. When we examined *vstm2b* expression by ISH at e12.5, we found no specific signal in lumbar DRG. By e13.5, however, we observed a number of clearly defined *vstm2b*^+^ cell bodies in lumbar DRG (Figure S3A). Moreover, *vstm2b* expression at e13.5 was restricted to DRG neurons co-expressing *trkC*, an early marker of pSNs and several other DRG sensory classes (Usoskin et al., 2014; Figure S3B), consistent with restriction of *vstm2b* expression to pSNs from the onset. Thus, *vstm2b* expression in lumbar pSNs begins ∼2 days after limb innervation, consistent with induction by a peripheral cue.

### Proprioceptor *vstm2b* expression influenced by dorsoventral character of limb mesenchyme

We next sought to characterize the involvement of individual limb tissues in patterning pSN *vstm2b* expression. We began by examining the contribution of limb mesenchyme, due to its established role in patterning the expression of muscle-type pSN genes *cdh13, sema5a*, and *crtac1*. pSN expression of *vstm2b* exhibits a clear bias along the dorsoventral limb axis, with nearly all dorsally innervating pSNs *vstm2b*^+^ and the majority of ventrally-innervating pSNs *vstm2b*^-^ (Figure 2G). We therefore assessed the role of dorsoventral mesenchymal character in patterning pSN *vstm2b* expression. We focused our analysis on pSNs supplying the shank, where the dorsoventral bias in pSN *vstm2b* expression is most pronounced: dorsal muscles (TA/EDL/PER) are innervated almost exclusively by *vstm2b*^+^ pSNs, and ventral shank muscles (GS/SOL/PLAN/FDL/FHL/TP) are supplied nearly entirely by *vstm2b*^-^ pSNs (Figures 2G and S2C).

We set out to characterize pSN *vstm2b* expression in mice with altered dorsoventral mesenchymal character. In wild type (WT) mice, expression of LIM-homeodomain transcription factor Lmx1b is restricted to dorsal mesenchyme, where it is necessary and sufficient to drive dorsal mesenchymal fate (Chen et al., 1998; Riddle et al., 1995; Vogel et al., 1995). We genetically manipulated mesenchymal dorsoventral character by driving ectopic expression of *lmx1b* in ventral limb mesenchyme. To achieve this, we crossed *Rosa*^*lsl*.*Lmx1b*^ mice to *Prx1*::*Cre* mice, resulting in Cre expression – and thus *lmx1b* expression – throughout limb mesenchyme beginning ∼e9.5 (cross termed *Prx1*^*Lmx1b*^; Figure 3D; Li et al., 2010; Poliak et al., 2016). Expression of *lmx1b* in ventral limb mesenchyme results in the conversion of muscle, bone and connective tissue to dorsal structures, yielding symmetric “double-dorsal” limbs (d/v → d/d’; Chen et al., 1998; Li et al., 2010; Poliak et al., 2016; Riddle et al., 1995; Vogel et al., 1995).

**Figure 3.**
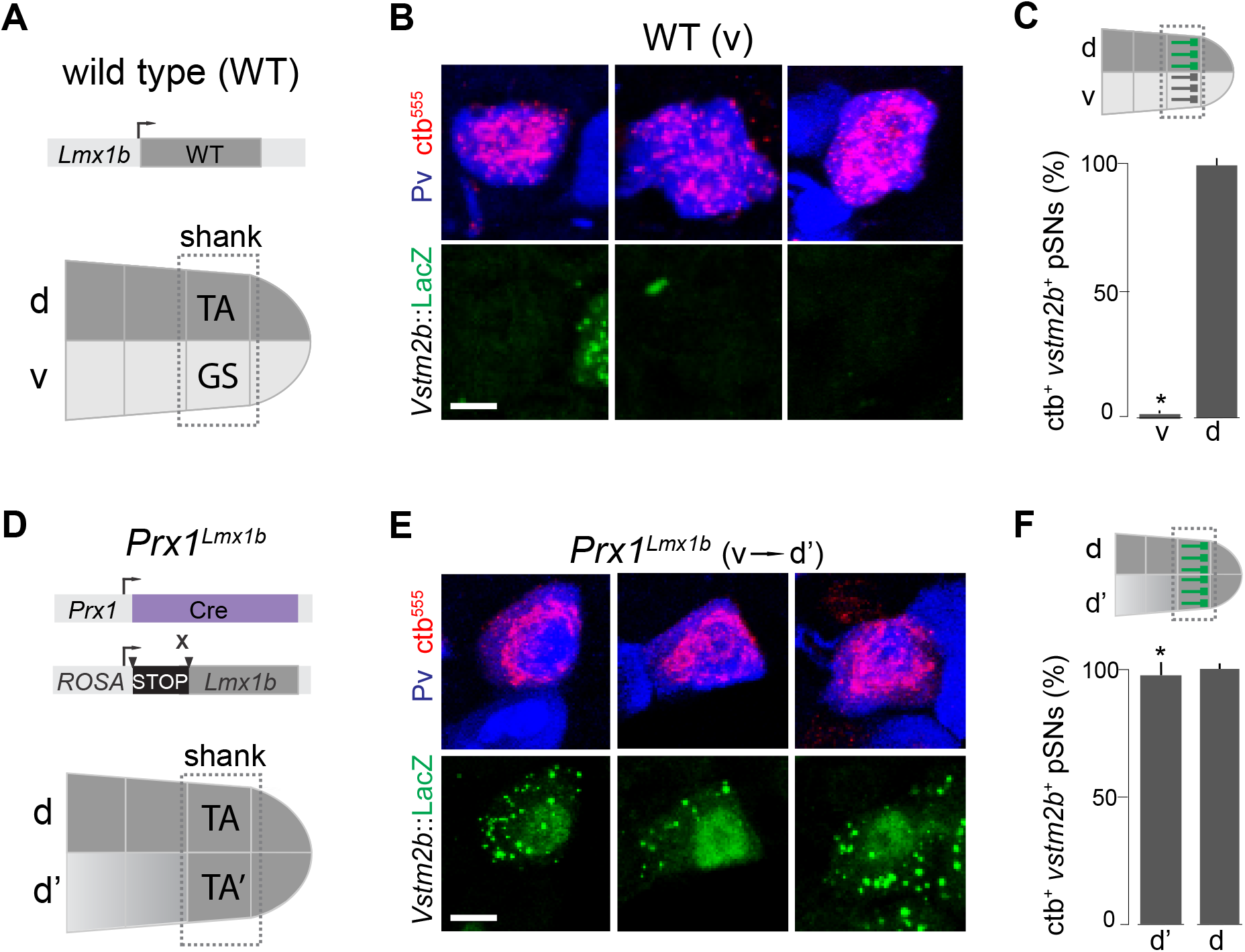
Dorsoventral Character of Limb Mesenchyme Influences pSN *vstm2b* Expression. (A-C) *vstm2b* expression in pSNs supplying dorsal (d) and ventral (v) shank muscles of wild type (WT) mice. (A) *Lmx1b* expression is restricted to dorsal limb mesenchyme in WT mice and is sufficient to instruct dorsal limb identity. (B) pSNs supplying ventral shank muscles (GS/Sol/Plan) were identified by ctb^555^ injection and Pv immunolabeling at P1. pSN *vstm2b* status was assessed by βgal immunolabelling in a *Vstm2b*::LacZ background. Scale bar, 10µm. (C) Fraction of WT ventral or dorsal (TA/EDL/Per) shank pSNs that express *vstm2b* (n=3 mice/injection). (D-F) *vstm2b* expression in pSNs supplying dorsal (d) and duplicated dorsal (d’) shank muscles of *Prx1*^*Lmx1b*^ mice. (D) Schematic depicting manipulation of limb mesenchymal identity. *Lmx1b* is expressed throughout the limb mesenchyme in *Prx1*::Cre, *Rosa*::lox-STOP-lox::*Lmx1b* mice (*Prx1*^*Lmx1b*^; arrowheads indicate *loxP* sites), imposing dorsal mesenchymal character (d’) on ventrally positioned limb domains. (E) pSNs supplying duplicated dorsal shank muscles (duplicated TA/EDL/Per) were assessed for *vstm2b* expression as described in (B). Scale bar, 10µm. (F) Fraction of dorsal or duplicated dorsal shank pSNs expressing *vstm2b* in *Prx1*^*Lmx1b*^ mice (n=3 mice/injection). (C, F) *, significant difference between WT ventral (v; GS/Sol/Plan) and *Prx1*^*Lmx1b*^ duplicated dorsal (d’; duplicated TA/EDL/Per) pSNs (p < 10^−5^, Student’s t test). No significant difference was detected between WT and *Prx1*^*Lmx1b*^ pSNs supplying dorsal limb domains. See also Figure S4.

In contrast to the drastic changes observed in muscle, bone and connective tissue patterning, generic aspects of pSN development - including the density of *pv*^+^ neurons in lumbar DRG and the number of MSs in shank muscles - are unchanged in *Prx1*^*Lmx1b*^ limbs (Poliak et al., 2016).

To facilitate our analysis of *vstm2b* induction, we generated a *Vstm2b*::LacZ reporter line (Figure S4A; Ringwald et al., 2011; see Supplemental Methods). We first assessed the pattern of *Vstm2b*::LacZ expression in lumbar DRG of heterozygotes. We observed that β-gal reporter was confined to Pv^+^, Rx3^+^ DRG neurons (Figure S4B), consistent with the endogenous distribution of transcript and protein. The proportion of β-gal^+^ pSNs in DRG L2-L6 (Figure S4C) was similar to proportion of *vstm2b*^+^ and Vstm2b^+^ neurons observed in these ganglia (see Figures 1D and 2B). To confirm the pSN pool specificity of LacZ reporter expression, we injected ctb^555^ into TA or GS muscles of P0 *Vstm2b*::LacZ heterozygotes and characterized the distribution of β-gal in Pv^+^ ctb^555^-labeled pSNs. At P1, we observed that 100% of backfilled TA and 0% of GS pSNs expressed β-gal (Figures S4D and S4E), thus demonstrating that the *Vstm2b*::LacZ allele recapitulates endogenous *vstm2b* expression.

We examined the profile of pSN *vstm2b* expression in *Prx1*^*Lmx1b*^ mice. To facilitate this analysis, we introduced the *Vstm2b*::LacZ allele into *Prx1*^*Lmx1b*^ animals. We examined retrogradely labeled pSNs for β-gal in these mice at P1, following injection of ctb^555^ into shank muscle groups at P0 (Figures 4B-4E). As expected, all pSNs supplying the endogenous dorsal shank muscles present in both *Prx1*^*Lmx1b*^ and WT mice expressed β-gal (Figures 4B-4E). However, while only ∼5% of Pv^+^ neurons WT supplying ventral shank muscles expressed β-gal, we found that ∼96% supplying ventrally positioned duplicated dorsal (d’) shank muscles were β-gal^+^ (Figures 4B-4E). We therefore conclude that the dorsoventral character of limb mesenchyme influences the pattern of pSN *vstm2b* expression.

**Figure 4.**
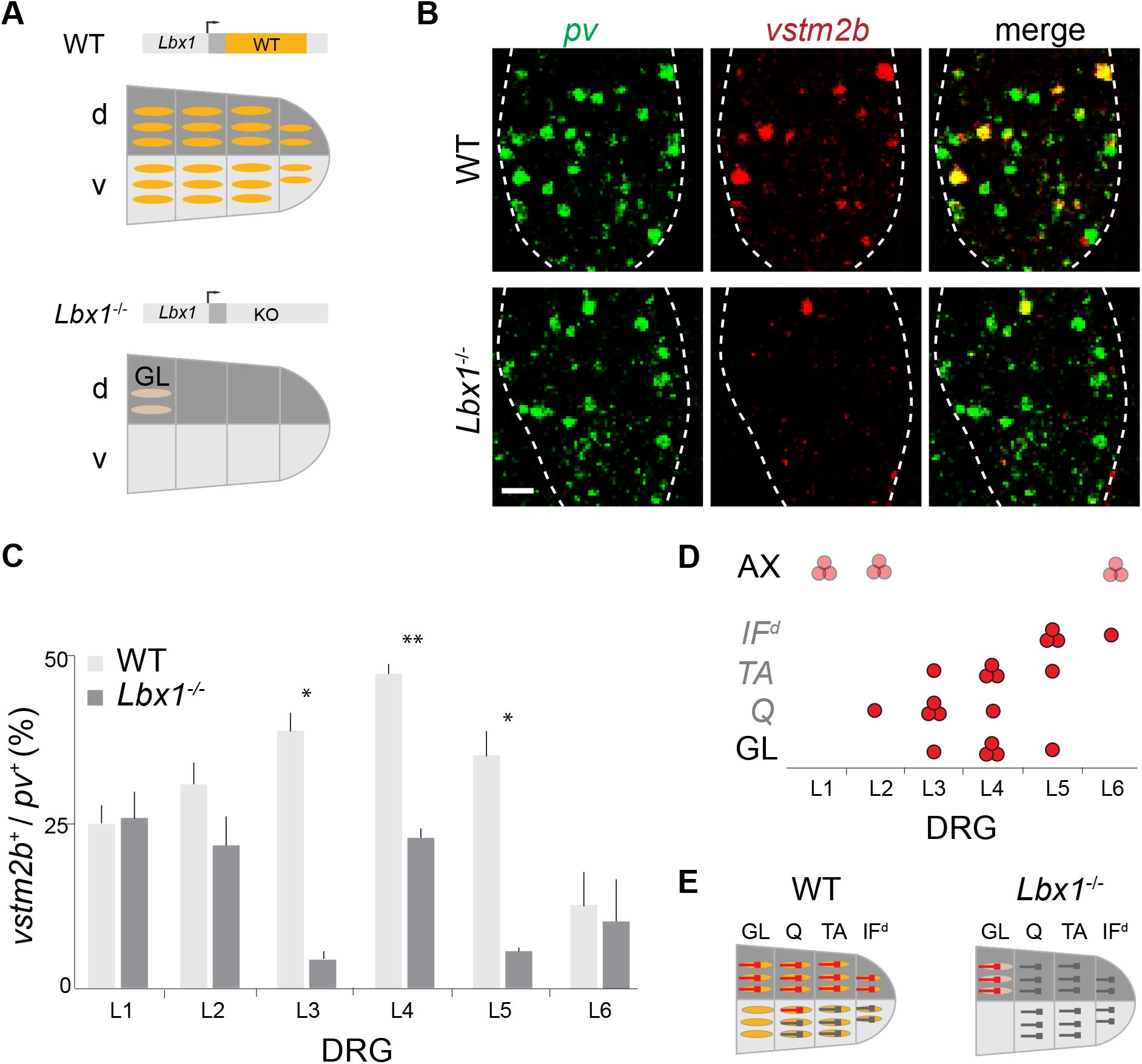
Limb Muscle is Required for pSN *vstm2b* Expression. (A) Schematic depicting manipulation of limb muscle. Top: in WT mice, myoblast expression of Lbx1 is required for their migration into the limb bud. Bottom: loss of Lbx1 results in inability of myoblasts to migrate and invade the limb bud, resulting in limbs devoid of most muscle. Only proximal hip musculature (GL, gluteus) persists in *Lbx1*^*-/-*^ mice. Orange: WT *Lbx1*^+^ muscle; tan: *Lbx1*^-/-^ muscle. (B-C) *vstm2b* expression in lumbar level pSNs of WT and *Lbx1*^-/-^ mice. (B) Double FISH for *pv* and *vstm2b* in e15.5 WT and *Lbx1*^-/-^ L5 DRG. Scale bar, 30µm. (C) Fraction of pSNs expressing *vstm2b* in lumbar DRG of e15.5 WT and *Lbx1*^-/-^ mice (n = 4 mice/genotype). Error bars represent SD. *p<0.001, **p<0.0001 (Student’s t test). (D) Schematic showing distribution of *vstm2b*^+^ pSN pools (red) in lumbar DRG of WT mice. AX: axial; IF^d^: dorsal intrinsic foot; TA: tibialis anterior; Q: quadriceps group; GL: gluteus group. Text in italic indicates loss of the muscle/muscle group in *Lbx1*^-/-^ mice. (E) Schematic comparing WT *vstm2b*^+^ pSN terminals to predicted *vstm2b* pSN pool status in *Lbx1*^-^/-mice. Red terminals: *vstm2b*^+^; grey terminals: *vstm2b*^-^.

### Limb muscle is required for proprioceptor *vstm2b* expression

In addition to patterning pSN gene expression, the dorsoventral character of limb mesenchyme instructs the pattern of limb muscle cleavage in dorsal and ventral limb compartments (Chen et al., 1998; Kardon et al., 2003; Poliak et al., 2016). We therefore questioned whether mesenchyme sets *vstm2b* expression directly by acting on pSNs or indirectly through its ability to pattern muscle.

To address this, we examined whether muscle is required to set the pattern of pSN *vstm2b* expression. We assessed pSN *vstm2b* status in *Lbx1*^-/-^ mice, in which myogenic precursor migration into the developing limb is inhibited such that hindlimbs are nearly devoid of skeletal muscle, with the exception in some embryos of gluteus (GL) muscle (Gross et al., 2000; Poliak et al., 2016; Figure 5A). Despite the absence of muscle, major peripheral nerve trajectories are preserved in *Lbx1*^-/-^ limbs, and *Lbx1*^-/-^ hindlimbs are invaded by DRG sensory axons (Phelan and Hollyday, 1990; Poliak et al., 2016). Moreover, the density of *pv*^+^ pSNs is unchanged in lumbar DRG at e15.5, shortly after limb innervation (Poliak et al., 2016).

**Figure 5.**
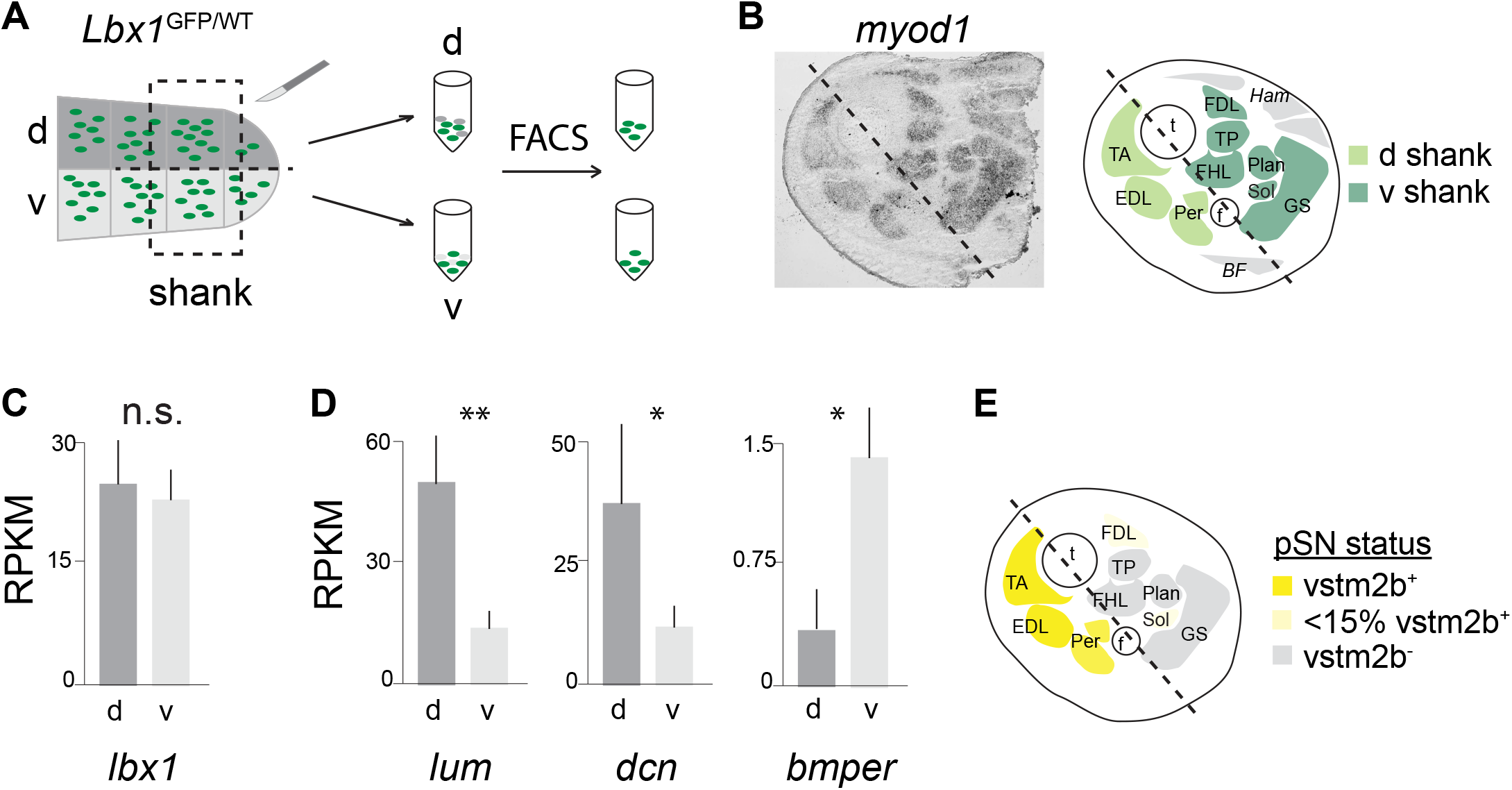
Gene Expression Profiling of Dorsal and Ventral Myoblasts. (A)Schematic depicting dorsal and ventral myoblast screen. Shanks of e13.5 *Lbx1*^GFP/WT^ mice were microdissected from the limb and cleaved along the dorsoventral axis. Dorsal and ventral shank samples were dissociated and FACS sorted to isolate GFP^+^ dorsal and ventral myoblasts. (B)ISH for *myod1* delineates individual shank muscles. The axis defined by the tibia (t) and fibula (f) cleanly separates dorsal (d) from ventral (v) myoblasts. (C-D) Gene expression levels in e13.5 dorsal (d) and ventral (v) shank myoblasts in reads per kilobase per million (RPKM). Error bars represent SD from three biological replicates. (C) Expression of generic myoblast gene *lbx1* does not differ significantly between d and v shank muscle samples. n.s. = not significant; p = 0.39. (D) Differentially expressed genes *lum* (FC = 3.69, p = 4.71×10^−9^), *dcn* (FC = 3.22, p = 3.4×10^−4^), and *bmper* (FC = 2.46, p =3.9×10^−3^). *, p<0.005; **, p<5×10^−9^. (E) Vstm2b expression pattern of shank pSNs. Shading of each muscle indicates the Vstm2b status of pSNs supplying it. See also Figure S5.

Postnatal lethality in *Lbx1*^*-/-*^ mice and the absence of limb muscle targets precluded examination of pSN pools defined by backfill. However, we reasoned that if *vstm2b* expression in hindlimb-innervating pSNs is dependent on muscle, we would see a reduction in the number of pSNs expressing *vstm2b* in lumbar DRG of *Lbx1*^-/-^ embryos.

We examined *vstm2b* expression in *pv*^+^ neurons at e15.5 in DRG L2-L6, thereby surveying all hindlimb-innervating pSNs, and observed a substantial reduction in the number of *vstm2b*^+^ pSNs in *Lbx1*^-/-^ versus WT littermate embryos (Figures 4B and 4C). Quantification revealed that the proportion of *vstm2b*^+^ pSNs decreased in all lumbar DRG, with significant differences between *Lbx1*^-/-^ and WT embryos observed for DRG L3-L5 (Figure 4C; L2: ∼31% pSNs *vstm2b*^+^ in WT versus ∼21% in *Lbx1*^-/-^, p = 0.08; L3: ∼39% WT versus ∼4% *Lbx1*^-/-^, p = 0.0002; L4: ∼47% WT versus ∼23% *Lbx1*^-/-^, p = 0.00001; L5: ∼35% WT versus ∼6% *Lbx1*^-/-^, p = 0.002; L6: ∼13% WT versus ∼10% *Lbx1*^-/-^, p = 0.39). In contrast, we found that the proportion of *vstm2b*^+^ pSNs in DRG L1, which contains only non-limb-innervating pSNs (de Nooij et al., 2013), was unchanged (Figure 4C; ∼25% WT versus ∼26% *Lbx1*^*-/-*^, p = 0.43).

What explains differences between lumbar DRG in the severity of *vstm2b*^+^ pSN loss observed in *Lbx1*^-/-^ animals? The distribution of pSN pools among lumbar DRG offers an explanation. A large fraction of pSNs in DRG L2 and L6 supply axial muscle (Figure 4D; de Nooij et al., 2013), which is unaffected in *Lbx1*^-/-^ embryos. Among limb-innervating pSNs, DRG L2 and L6 house pSNs supplying Q and IF muscles, respectively (Figure 4D; unpublished observation). The relatively minor decrease in *vstm2b*^+^ pSNs in DRG L2 and L6 can be attributed to loss of *vstm2b* expression by Q and IF pSNs, whose targets are lost in *Lbx1*^-/-^ embryos (Figures 4C-E). In DRG L3-L5, pSNs almost exclusively supply hindlimb muscles, many of which receive *vstm2b*^+^ pSN innervation in WT animals (Figures 1D and 2B). Thus, larger fractions of *vstm2b*^+^ pSNs are lost in these DRG in *Lbx1*^-/-^ embryos due to the absence of target limb muscle (Figures 4C-E). Residual *vstm2b*^+^ pSNs in these ganglia likely supply GL group muscles, which persist in *Lbx1*^-/-^ embryos (Gross et al., 2000). GL group pSNs reside in DRG L3-L5, with the majority located in L4 (Figure 4D; unpublished observation), consistent with the distribution of remaining *vstm2b*^+^ pSNs in these ganglia in *Lbx1*^-/-^ mice (Figures 4C and 4E).

Taken together, these data demonstrate that the presence of limb muscle is required for induction of the normal pattern of pSN *vstm2b* expression.

### Embryonic muscles possess distinct dorsoventral identities

Our finding that muscle is required for normal induction of pSN *vstm2b* expression suggests that limb muscle may act as a source of patterning cue. If muscle is acting in an instructive capacity, muscles must differ in their molecular makeup sufficiently early in embryonic development to direct the specification of pSNs. The near restriction of *vstm2b* expression to pSNs supplying dorsal hindlimb by e13.5 implies that dorsal and ventral muscles must exhibit distinct molecular identities around this time point.

To explore whether muscle exhibits early signatures of dorsoventral identity that might instruct pSN *vstm2b* expression, we compared the gene expression profiles of dorsal and ventral hindlimb myoblasts. We limited our analysis to shank myoblasts due to the near complete restriction of *vstm2b* expression to pSNs supplying dorsal muscle at this limb level (Figures 2G and 5E). Limb myoblasts are marked by expression of *lbx1* (Dietrich et al., 1998; Jagla et al., 1995). We utilized *Lbx1*::GFP heterozygotes (*Lbx1*^GFP/WT^; Figure 5A), in which myoblasts can be distinguished from other limb tissues by GFP expression (Gross et al., 2000), to isolate myoblasts from surrounding tissues. We isolated shank segments from *Lbx1*^GFP/WT^ embryos at e13.5, when *vstm2b* expression is first observed in lumbar pSNs and prior to the elongation of myoblasts into multinucleate muscle fibers. We then separated dorsal from ventral shank by cleaving the segment along the axis defined by the tibia and fibula (Figures 5A and 5B). After dissociation of dorsal and ventral samples, we separated GFP^+^ myoblasts from GFP^-^ lineages by FACS (Figures 5A, S5A and S5B). Finally, total RNA was extracted and cDNA libraries were prepared for RNA-sequencing (RNA-seq) analysis.

We first assessed the composition of dorsal and ventral samples by examining expression of myoblast marker *lbx1*. Our RNA-seq results did not show a significant difference in *lbx1* expression between dorsal and ventral samples (p=0.39; Figure 5C), indicating an equal representation of myoblasts in our dorsal and ventral cDNA libraries.

We next searched for genes differentially expressed by dorsal and ventral shank myoblasts. Analysis of our RNA-Seq data yielded 105 and 127 genes enriched >2-fold in dorsal and ventral myoblasts, respectively (p<0.05). We narrowed our focus to genes positioned to directly assign pSN *vstm2b* expression – namely, transmembrane and secreted proteins and molecules with known functions as ligands in signaling pathways – thereby reducing the number of candidates to 23 dorsal and 18 ventral genes (Table S1). In situ hybridization identified 3 genes encoding secreted modulators of cell signaling pathways with clear qualitative expression differences between dorsal and ventral shank muscles: dorsally-enriched *lumican* (*lum*) and *decorin* (*dcn*), and ventrally-enriched *BMP binding endothelial regulator* (*bmper*) (*lum*: fold change FC = 3.7, p = 4.71×10^−9^; *dcn*: FC = 3.22, p = 3.4×10^−4^; *bmper*: FC = 2.46, p = 3.9×10^−3^; Figures 5D and 6, Figures S6A-S6C).

To obtain quantitative expression profiles of dorsal- and ventral-biased genes in individual muscles, we performed RNAScope fluorescent in situ hybridization on shanks of WT mice. We quantified the expression of *dcn, lum*, and *bmper* in shank muscles, whose boundaries we defined based on *myod1* expression (Figure S6) at e14.5, when the pattern of hindlimb muscle cleavage is well defined (Lance-Jones, 1979) For dorsal muscle genes *lum* and *dcn*, we observed high expression in TA muscle and weaker expression within GS muscle (Figures 6A-6F; *lum*: 7% and *dcn*: 19% TA expression level), consistent with upregulation in dorsal myoblasts predicted by our RNA-Seq screen. In contrast, *bmper* signal was highly enriched in dorsal GS muscle compared to ventral TA (Figures 6G-6I; 7% GS expression level), consistent with enrichment in ventral myoblasts.

**Figure 6.**
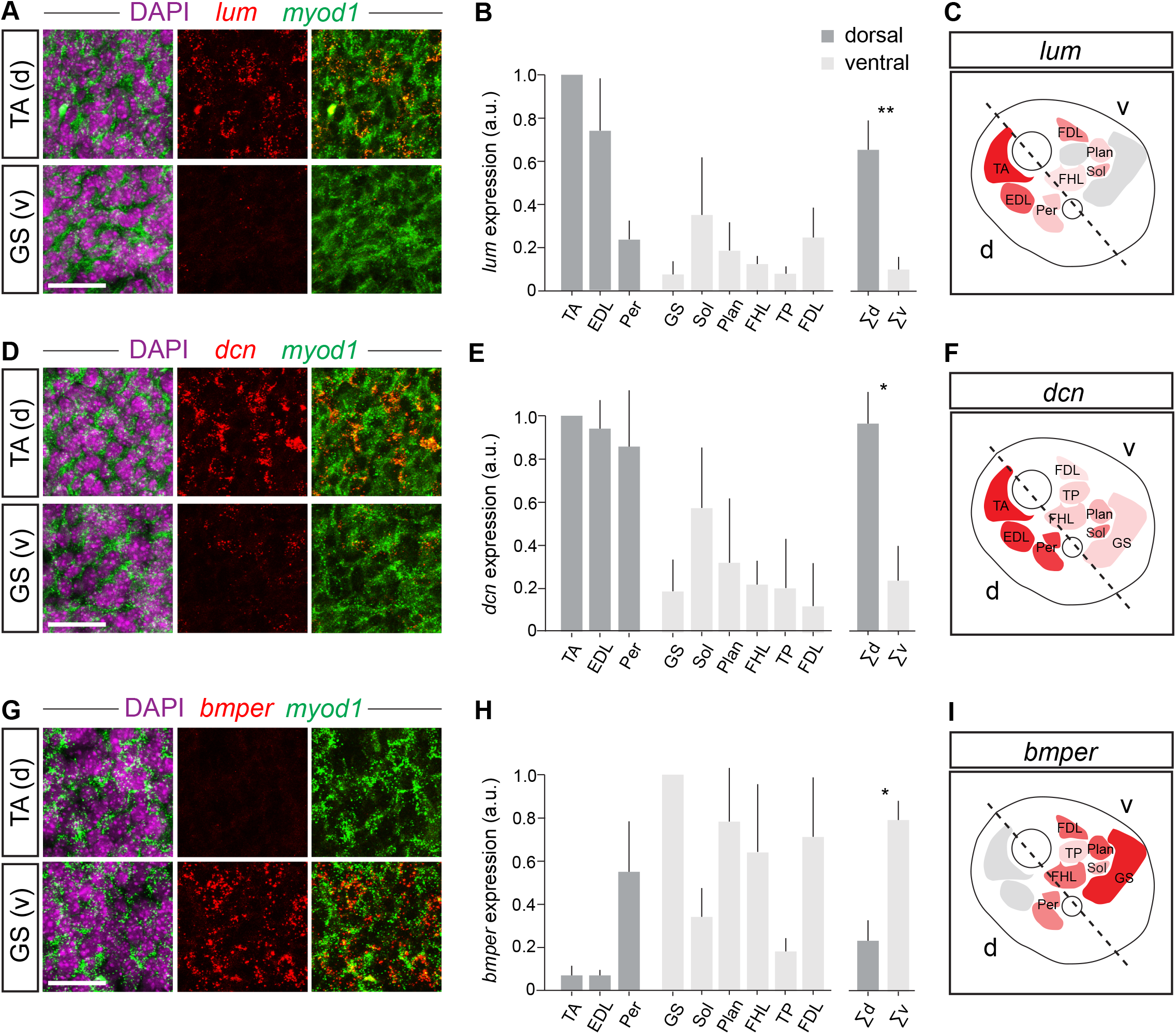
Early Molecular Distinctions between Dorsal and Ventral Muscles. (A-I) RNAScope *in situ* hybridization in e14.5 shank shows differential expression of *lum* (A-C), *dcn* (D-F), and *bmper* (G-I) in individual dorsal (d) and ventral (v) muscles. (A, D, and G) RNAScope shows TA-or GS-biased expression of *lum* (A), *dcn* (D), and *bmper* (G) in muscle cells defined by DAPI and *myod1* expression. Scale bars, 30µm. (B, E, and H) Normalized expression levels of *lum* (B), *dcn* (E), and *bmper* (H) in dorsal (d; TA, EDL, Per) and ventral (v; GS, Sol, Plan, FHL, TP, FDL) shank muscles. Error bars represent SD. *p < 0.05, **p < 0.005 (paired Student’s t test). (C, F, and I) Shank cross-sectional schematics indicate relative gene expression level in each shank muscle. Color intensity is scaled relative to TA (*lum, dcn*) or GS (*bmper*) muscle (*lum*: n=7; *dcn*: n=4; *bmper*: n=7). See also Figure S6.

We expanded this analysis to all dorsal and ventral shank muscles. Relative to TA expression (100%), *lum* was expressed at high levels in dorsal EDL (74%) muscle, at intermediate levels in dorsal Per (24%) and ventral Sol (35%), Plan (19%), FDL (25%), and FHL (12%) muscles, and was nearly absent from ventral TP (7%; Figures 6B-6C and S6A). Likewise, *dcn* was highly upregulated in all dorsal muscles (EDL: 93%, Per: 86%) and downregulated in all ventral muscles (Figures 6E-6F and S6B; Sol: 57%, Plan: 32%, FHL: 22%, TP: 20%, FDL: 12%). Finally, relative to GS expression (100%), we found that *bmper* was expressed highly in ventral Plan (78%), FHL (64%), and FDL (71%) muscles and at lower levels in ventral TP (18%) and Sol (34%). In dorsal muscles, *bmper* was expressed at an intermediate level in the Per group (55%), and was nearly absent from EDL muscle (7%; Figures 6H-6I and S6C).

Taken together, these results demonstrate that dorsal and ventral muscles exhibit distinct profiles of gene expression early in embryonic development, consistent with an instructive role for muscle in setting the pattern of pSN *vstm2b* expression.

## Discussion

Neither the extent of pSN pool identities nor the mechanisms that establish them are fully understood. We describe preferential expression of *vstm2b* by pSNs supplying dorsal muscle within the hindlimb. The profile of pSN *vstm2b* expression is altered in the absence of target limb muscle, demonstrating a requirement for myoblasts in assigning aspects of pSN pool identity. In line with our finding that muscle instructs pSN gene expression in neurons supplying dorsal muscle, we find that dorsal and ventral myoblasts exhibit molecular distinctions at the time of pSN termination in the limb bud. These data reveal a previously uncharacterized feature of muscle differentiation and identify candidate patterning molecules aligned with the pSN profile of *vstm2b* expression.

### Vstm2b in the context of sensory-motor circuits

Previous studies of pSN diversity have identified molecular markers of defined pSN subpopulations (Chen et al., 2002; Fukuhara et al., 2013; de Nooij et al., 2013; Poliak et al., 2016). Notably, expression of pSN genes *cdh13, sema5a*, and *crtac1* defines pSN pools supplying dorsal or ventral domains of the hindlimb, linking pSN gene expression to domain of limb innervation (Poliak et al., 2016). However, these markers define only pSNs that supply distal hindlimb domains, suggesting that our understanding of pSN pool organization – and by extension, its specification – is incomplete. Muscle-by-muscle analysis of pSN gene *vstm2b* expression reveals expression in neurons innervating dorsal muscles along the proximodistal extent of the hindlimb, extending our understanding of hierarchical pSN pool organization by identifying a sensory correlate of motor neuron divisional identity.

Could Vstm2b be involved in establishing specific connections with motor neurons? Dorsoventral distinctions in pSN gene expression are likely to instruct mediolateral targeting in the spinal cord, since motor neurons supplying dorsal versus ventral limb muscle targets reside in the lateral and medial divisions of the LMC, respectively (Sürmeli et al., 2011). Accordingly, a shared molecular identity among pSNs supplying dorsal muscles might influence the mediolateral choice of pSN axons. The transmembrane Ig-domain structure of Vstm2b supports a function in axon guidance or cellular recognition. Vstm2b lacks an intracellular domain, so if it is to regulate synaptic specificity, it will likely bind receptors in both cis and trans. We noted *vstm2b* expression in spinal motor neurons and interneurons, although the subtype specificity of its expression in these populations – if any – has not been characterized. Defining motor pool expression of *vstm2b* might therefore provide insight into the mode of target recognition by *vstm2b*-expressing afferents.

### Induction of *vstm2b* by peripheral cues

*Vstm2b* differs from previously identified pSN pool markers in two respects. First, whereas *cdh13, sema5a*, and *crtac1* are expressed by DRG cutaneous sensory neurons in addition to pSNs, *vstm2b* expression in DRG is restricted to pSNs. Second, *vstm2b* induction is delayed with respect to the onset of *cdh13* expression by ∼1 day. Both differences are consistent with our finding that muscle is required for the induction of *vstm2b* expression in pSNs. *Cdh13, sema5a*, and *crtac1* are induced in a muscle-independent manner by mesenchymal cues, which exist in diffuse gradients along the proximodistal and dorsoventral limb axes (Bénazet and Zeller, 2009). In contrast, muscle-derived cues are likely more localized. Thus, while both pSN and cutaneous axons are exposed to limb mesenchymal cues as they project toward their targets, cutaneous sensory neurons might avoid exposure to local muscle-derived signals. Moreover, differences in the onset of *cdh13* and *vstm2b* expression can be explained by the timing of exposure to each tissue: pSN axons traverse mesenchyme, and are thus exposed to mesenchymal cues, prior to exposure to target muscle-derived cues.

The emergence of pSN *vtsm2b* expression appears linked to the positional specification of limb tissues. In contrast to pSN pool marker *cdh13*, which is induced directly by cues from limb mesenchyme in a muscle-independent manner, we find that pSN *vstm2b* expression is both influenced by limb mesenchyme and requires the presence of limb muscle. How might both tissues be involved in inducing *vstm2b* expression?

Limb muscle fate appears to be extrinsically specified by cues deriving from limb mesenchyme (Chevallier et al., 1977; Jacob and Christ, 1980; Kardon et al., 2003). Indeed, the morphology of limb muscle is influenced by the dorsoventral character of limb mesenchyme (Chen et al., 1998; Poliak et al., 2016; Riddle et al., 1995; Vogel et al., 1995). One model of *vstm2b* induction therefore holds that the effects of mesenchyme and muscle are sequential: limb mesenchyme patterns the dorsoventral molecular identity of muscle, which in turn specifies dorsoventrally restricted gene expression in pSNs. Alternatively, mesenchyme and muscle might act in parallel to supply multiple patterning cues that converge to drive *vstm2b* expression in subsets of pSNs.

### Early muscle positional identities align with proprioceptor *vstm2b* expression

The sequential model of mesenchymal and muscle action, in which muscle plays an instructive role, requires spatially restricted gene expression in limb muscle that aligns with the pattern of pSN *vstm2b* expression. Our screen demonstrates the distinct molecular character of dorsal and ventral shank muscles, exemplified by *dcn, lum*, and *bmper* expression, and defines molecular differences that align both spatially and temporally with the *vstm2b* induction in pSNs. Despite the known influence of dorsoventral position on muscle differentiation, dorsal and ventral myoblasts have never been systematically profiled for early molecular signatures of subtype identity; to our knowledge, genes that distinguish dorsal and ventral muscles this early in development - shortly after cleavage and around the time of innervation by pSNs - have not been identified previously.

The proteins encoded by these genes are secreted modulators of cell signaling. *Dcn* and *lum*, both expressed in dorsal shank muscle, are small leucine-rich proteoglycans that modulate a number of cell signaling pathways, including BMP and TGFβ (Schaefer and Iozzo, 2008). Consistent with mesenchymal control of spatially restricted muscle gene expression, *dcn* and *lum* are downregulated in ventralized limbs of *lmx1b*^-/-^ embryos (Gu and Kania, 2010; Krawchuk and Kania, 2008). Alternatively, ventrally-enriched *bmper* could set the pattern of *vstm2b* expression by locally modulating BMP signaling in ventral limb, thereby restricting *vstm2b* expression to proprioceptors supplying dorsal muscle through a repressive mechanism. Indeed, mesenchymally-derived BMPs are distributed throughout the limb bud, making this signaling pathway an attractive candidate (Norrie et al., 2014; Pizette et al., 2001). Future studies will clarify the molecular mechanisms by which spatially restricted cues from multiple tissue sources are integrated to generate the neuronal diversity required to meet the biomechanical demands of the limb.

## Experimental Procedures

### Experimental Model and Subject Details

*Prx1::*Lmx1b (*Prx1::*Cre; *Rosa26::lox-STOP-lox*.Lmx1b; Li et al., 2010; Logan et al., 2002; Poliak et al., 2016) and *Lbx1*^*-/-*^ (Gross et al., 2000; Poliak et al., 2016) mouse strains have been described previously. BALB/cJ mice were used as the wild type strain. Both male and female mice were used for all experiments based on availability and were maintained under standard husbandry and housing conditions. All experiments were performed in accordance with the National Institutes of Health Guidelines on the Care and Use of Animals and were approved by the Columbia University animal care and use committee. Generation of *Vstm2b*::LacZ mice is described in Supplementary Methods.

## Method Details

### Retrograde labeling of muscle-specific proprioceptors

Muscles of anesthetized p0 mice were pressure injected with 1% ctb^555^ (Life Technologies) using a hand-pulled capillary and aspirator tube. Mice were sacrificed by decapitation at p1 and eviscerated, then lumbar spinal cords were fresh-frozen for *in situ* hybridization or subjected to ventral laminectomy and fixed for immunohistochemistry as described below. The specificity of muscle injections was confirmed by muscle dissection and examination under a fluorescence microscope, as well as by the position and clustering of retrogradely labeled spinal motor neurons in laminectomized animals. Animals with injections restricted to the desired muscle or muscle group were processed for further analysis.

### In situ hybridization

Chromogenic *in situ* hybridization was performed on 12-18 µm cryostat sections using digoxygenin (DIG)-labeled probes as previously described (Schaeren-Wiemers and Gerfin-Moser, 1993). Fluorescence in situ hybridization (FISH) was performed on 12-16 µm cryostat sections hybridized with DIG- and fluorescein isothiocyanate (FITC)-labeled probes using the FITC/Cy5 TSA Plus Fluorescence System for signal amplification according to the manufacturer’s instructions. All in situs were performed on fresh-frozen tissue.

Antisense probes were generated by polymerase chain reaction (PCR) from mouse p1 lumbar DRG or e13.5 whole embryo cDNA libraries. cDNA libraries were synthesized using the SuperScript III First-Strand Synthesis System for RT-PCR (Invitrogen) from RNA purified using the Absolutely RNA Miniprep Kit (Agilent). Primers were sourced from the Allen Mouse Brain Atlas (Lein et al., 2007; see Table S2). Reverse primers were appended with the T7 polymerase binding sequence to enable direct *in vitro* transcription. PCR products were gel purified, sequenced and used as template for transcription of antisense probes with DIG or FITC RNA labeling kits (Roche).

### Immunohistochemistry

Tissue used for immunohistochemistry was fixed for 2 h or overnight in 4% PFA/0.1M PB at 4C. Following PBS washes, tissue was dehydrated in 30% sucrose/0.1M PB and embedded in OCT (Tissue Tek) for cryosectioning (10-16 µm). With the exception of spindle labeling experiments, tissue sections were incubated overnight at 4C with primary antibodies diluted in 0.1% Triton X-100/PBS solution (0.1% PBT). Following 3x PBS washes, sections were incubated with secondary antibodies diluted in 0.1% PBT for 1 h at room temperature.

For spindle labeling, p3-5 hindlimbs with skin removed were fixed in 4% PFA/0.1M PB for 2 h at 4C and dehydrated in 30% sucrose/0.1M PB. Individual muscles were dissected and embedded in OCT for cryosectioning (25 µm). Sections were submerged in 0.3% PBT primary antibody solution and incubated for 48 h at 4C.

The following antibodies were used in this study: chicken anti-Pv (1:10000, de Nooij et al., 2013), rabbit anti-Rx3 (1:50000, Kramer et al., 2006), chicken anti-βgal (1:5000, Abcam ab9361), and rabbit anti-vGluT1 (1:16000, de Nooij et al., 2013). Guinea pig anti-Vstm2b was generated against purified His-tagged extracellular region of mouse Vstm2b protein (cell body: 1:64000, muscle spindle: 1:8000; see Supplemental Methods). Secondary antibodies were generated in donkey and conjugated to FITC, Cy3 or Cy5 (FITC and Cy5: 1:500, Cy3: 1:1000; Jackson Immunoresearch Laboratories).

### RNA-Sequencing of dorsal and ventral muscle cells

Prior to sample collection, e13.5 embryos were rapidly genotyped under a fluorescence microscope to identify animals carrying the *Lbx1*^GFP^ allele. Dorsal and ventral shank segments were isolated using a microdissection knife (Sharpoint Microsurgical Knife, 25 gauge). The shank was isolated by cleaving just proximal to the knee and distal to the ankle, then dorsal and ventral shank were separated by cleaving along the visible plane of the tibia and fibula. Cell suspensions were prepared from dorsal and ventral shank segments as described previously (Campbell et al., 2012) for purification of GFP^+^ myoblasts by flow cytometry (MoFlo Astrios EQ, Beckman Coulter). Each biological replicate contained both left and right shank segments from all *Lbx1*^GFP/WT^ embryos in one litter (i.e., 4-6 mice per replicate).

Total RNA was extracted using the RNeasy Plus Mini Kit (Qiagen) according to the manufacturer’s protocol. cDNA libraries were constructed using the TruSeq RNA prep kit (Illumina) according to the manufacturer’s instructions and sequenced on the Illumina HiSeq 2500 platform to a depth of 25-30 million single-end 100 bp reads per sample at the Columbia Genome Center.

### RNA-Scope Assay

Briefly, e14.5 embryos were decapitated and fixed overnight in 4% PFA/0.1M PB at 4C. Following dehydration in 30% sucrose, shanks were embedded in OCT for cryosectioning (16µm). RNA-Scope transcript detection was performed using the V1 kit according to the manufacturer’s protocol with probes purchased from Advanced Cell Diagnostics (see Table S2).

## Imaging

Images were collected using Zeiss 880, 710 and 510 laser scanning confocal microscopes.

## Quantification and Statistical Analysis

RNAScope hindlimb expression data was quantified in ImageJ. *Myod1* signal was used to define each muscle as an independent region of interest (ROI). ROI masks were applied to the candidate gene image, and the area covered by candidate gene signal was computed for each ROI using the Analyze Particles function. A minimum of three sections per animal were selected from multiple points along the proximodistal axis of the shank for analysis. All images for a given candidate gene were acquired using identical laser power and gain configurations.

## Author Contributions

A.L.N performed all experiments and data analysis. S.B.M. generated critical reagents. A.L.N. and T.M.J. devised the project and designed experiments. A.L.N. prepared the manuscript.

## Acknowledgements

We are grateful to Qiaolin Liu, David Wu and Brenda Abdulmessih for technical assistance. Ira Schieren performed FACS, and Monica Mendelsohn and Nataliya Zabello generated *Vstm2b::*LacZ mice. *Lbx1* knockout mice were the generous gift of Martyn Goulding. We thank Jane Dodd and Andrés Bendesky for comments on the manuscript, and Sebastian Poliak and Joriene de Nooij for useful discussions. We are grateful to Barbara Han, Erica Famojure, Myles Marshall, and Kathy MacArthur for administrative assistance. A.L.N. was supported by an NIH training grant. T.M.J. was supported by the Howard Hughes Medical Institute and grants from NINDS, The Leila and Harold Mathers Foundation, Project ALS, and the Simons Foundation.

