## Supplement for "Muscle-derived Cues are Required to Specify Proprioceptor Pool Identity"

### Supplemental Experimental Procedures

**RNA-Seq screen of TA and GS pSNs** (Previously reported in Poliak et al., 2016)

TA or GS muscles of p0 *Pv*::YFP pups (*Pv*::Cre, *Thy1*::lox-STOP-lox::YFP mice; Buffelli et al., 2003; Hippenmeyer et al., 2005) were injected with cholera toxin B sub-unit/Alexa$^{555}$ (ctb$^{555}$; 1% dilution in PBS; Life Technologies). The following day, DRG containing ctb$^{555}$-labeled neurons were removed and dissociated (Malin et al., 2007), and individual YFP$^{+}$, ctb$^{555+}$ proprioceptors were identified and purified by aspiration (Hempel et al., 2007). Total RNA was extracted from three samples of TA and GS proprioceptors, each containing 25-30 neurons, using a PicoPure RNA isolation kit (Arcturus). cDNA was synthesized using the Ovation RNA-Seq System V2 kit (Nugen). cDNA libraries were constructed using the Nextera DNA Sample Prep kit (Illumina) and sequenced on an Illumina HiSeq 2000 to a depth of 25-30 million paired-end reads per sample. Analysis was carried our using ExpressionPlot software (Friedman and Maniatis, 2011). Briefly, reads were aligned to the mouse mm9 genome assembly and annotated using the Ensembl v62 database. Differential expression analysis was performed using DESeq. Data were deposited into the GEO repository under the accession number GSE71028.

**Generation and validation of *Vstm2b*::LacZ Mice**

*Vstm2b*::LacZ mice were generated from targeted ES cells obtained from the EUCOMM division of the International Knockout Mouse Consortium project (Ringwald et al., 2011; Skarnes et al., 2011). Chimeras were produced by Monica Mendelsohn and Nataliya Zabello at Columbia University by injecting BALB/cJ morulae with *vstm2b* targeted ES cells (clone F02). Offspring of chimeras carrying the targeted allele were bred to *Protamine*::Cre mice (O'Gorman et al., 1997) to achieve male germline deletion of the SV40-Neo selection cassette, yielding a *Vstm2b*::LacZ reporter allele (Figure S4A).

**Generation of Vstm2b Polyclonal Antibody**

Sequence encoding a segment of the Vstm2b extracellular domain (aa129-144) was cloned into pQE32 vector containing a 6xHis tag via BamHI and SalI sites. Vstm2b protein production was induced with IPTG. Bacterial pellet was denatured with guanidine-HCl and insoluble protein was removed by centrifugation. Vstm2b-His was purified from the supernatant on a column containing NiNTA resin (Biorad dispo column) and eluted with imidiazole buffer. Soluble protein in PBS was used to immunize guinea pigs (Covance). Following immunization, bleeds were collected at regular intervals and tested on tissue sections for immunoreactivity. Serum was collected once specific reactivity was observed.

**RNAScope Data Quantification**

Images (20x tiled, one z plane) were processed and quantified in ImageJ. Individual shank muscles were defined as regions of interest (ROIs) using *myod1* labeling as a guide, taking care to exclude autofluorescent vasculature. The channel containing gene of interest (GOI) data was manually thresholded using values determined from control sections not treated with probe for the GOI. The image was binarized and the Analyze Particles function was used to quantify the area within the ROI covered by GOI puncta. The obtained GOI area value was normalized by dividing by the ROI area of the muscle. This value was then divided by TA (*lum*, *dcn*) or GS (*bmper*) value to obtain the relative GOI expression level for each muscle (Figure 6). A minimum of 3 sections per animal collected at different points along the proximodistal axis were used.

### Supplemental Figure Legends

**Figure S1. RNA-Sequencing Screen of TA and GS pSNs,** Related to Figure 1

Schematic depicting workflow of screen for gene expression differences in tibialis anterior (TA) and gastrocnemius (GS) pSNs. Pv: parvalbumin; YFP: yellow fluorescent proten; $ctb^{555}$: cholera toxin subunit b conjugated to Alexa 555. This screen was previously published in Poliak et al., 2016.

**Figure S2. Vstm2b Expression in Muscle-Type Proprioceptors**, Related to Figure 2

(A) Fraction of ($Pv^{+}$ $Rx3^{-}$), ($Pv^{-}$ $Rx3^{+}$), and ($Pv^{-}$ $Rx3^{-}$) DRG neurons marked by Vstm2b immunolabeling in DRG L1-6 at P3. n=3 mice.

(B) $vGluT1^{+}$ Golgi tendon organ (GTO) afferents are marked by Vstm2b immunolabeling in TA but not GS muscle at P5. Scale bar, 50μm.

(C) Fraction of $vGluT1^{+}$ muscle spindle (MS) afferents marked by Vstm2b immunolabeling in hip, thigh, shank and foot hindlimb muscles at P3-5.

(D) Comparison of the fraction of *vstm2b*$^{+}$ muscle-type pSNs detected by fluorescent in situ hybridization (FISH) in backfilled $ctb^{+}$ cell bodies vs. $Vstm2b^{+}$ immunolabeled MSs in dissected muscles. Σd thigh = Q (VL, VM, RF+VI); Σv thigh = H/A (ST, Grac, BF, SM, Add). n.s., $p > 0.05$ by Student's t test.

**Figure S3. *Vstm2b* Expression Onset**, Related to Figures 3 and 4

(A) *vstm2b* is observed in lumbar DRG beginning at e13.5. Left: *vstm2b* ISH at e12.5. Image is intentionally overexposed such that background signal can be used to define boundaries of spinal cord and DRG. Right: *vstm2b* ISH at e13.5. *vstm2b* signal is clearly observed in spinal motor neurons and a subset of DRG neurons.

(B) Fluorescent in situ hybridization (FISH) reveals restriction of *vstm2b* expression to *trkC*$^{+}$ neurons in e13.5 lumbar DRG.

**Figure S4. *Vstm2b*::LacZ Allele Recapitulates Endogenous Vstm2b Expression**, Related to Figure 3

(A) Top: ES cells carrying the targeted locus shown were obtained from the EUCOMM project (Skarnes et al 2011). Bottom: A *Protamine*::Cre mice allele was used to excise the neo cassette and exons 2-4, resulting in a *Vstm2b*::LacZ allele.

(B-C) β-gal protein from the *Vstm2b*::LacZ locus is restricted to $Pv^+$ $Rx3^+$ pSNs. (B) Immunolabeling in L4 DRG at P3. pSNs are defined by the intersection of Pv and Rx3 labeling. Scale bar, 10μm. (C) Quantification of β-$gal^+$ pSNs at P3 in lumbar level DRG (L, lumbar level; n=3; >100 neurons/level). Error bars represent SD.

(D-E) Vstm2b status of TA and GS pSN pools. (D) TA and GS pSNs identified by $ctb^{555}$ and Pv immunolabeling at P3. Arrowheads indicate TA or GS pSNs, as defined by the coincidence of Pv immunolabeling and $ctb^{555}$. Closed: $Vstm2b^+$ pSN; open: $Vstm2b^-$ pSN. (E) Fraction of TA and GS pSNs marked by Vstm2b immunolabeling (TA: n=4, GS: n=5). Error bars represent SD. *, significant difference between TA and GS samples (Student's t test, $p < 10^{-5}$). Scale bar, 10μm.

**Figure S5. Dorsal versus Ventral Myoblast Screen,** Related to Figure 5

(A) Representative FACS of myoblast samples. Viable myoblasts were gated by $GFP^+$ $DAPI^+$ status. Collected $GFP^+$ $DAPI^+$ cells represented ~0.05% of all dissociated cells. Dorsal (d) and ventral (v) samples did not differ significantly in composition.

(B) Volcano plot showing statistical significance (p-value) versus fold-change of genes expressed in dorsal and ventral myoblast samples. Horizontal line indicates p-value cutoff of 0.05; vertical lines indicate fold change cutoff of 2. Red: genes with confirmed muscle-specific expression profiles; green: generic myoblast marker.

**Figure S6. Differential Gene Expression in Shank Muscles**, Related to Figures 5 and 6

Expression profiles of *lum* (A), *dcn* (B), and *bmper* (C) in transverse sections of e14.5 WT shank as detected by RNAScope in situ hybridization. Muscle is defined by *myod1* expression and mesenchyme by *prx1* expression. t, tibia; f, fibula. Scale bars, 200μm. Dorsal muscle: TA (tibialis anterior), EDL (extensor digitorum longus), Per (peroneal group). Ventral muscle: GS (gastrocnemius), Sol (soleus), Plan (plantaris), FHL (flexor hallucis longus), TP (tibialis posterior); * indicates position of FDL (flexor digitorum longus) in more proximal sections.

### Supplemental Tables

A

| Gene | | FC d vs. v | p-value |
|---|---|---|---|
| Ms4a4d | membrane-spanning 4-domains, subfamily A, member 4D | 33.6136639 | 0.009863255 |
| Epyc | epiphycan | 31.39561164 | 0.013433386 |
| Clec12b | C-type lectin domain family 12, member B | 26.71193276 | 0.041313072 |
| Tnfsf18 | tumor necrosis factor (ligand) superfamily, member 1 | 20.69182464 | 0.015381066 |
| Kera | keratocan | 9.329802949 | 1.18E-38 |
| Cntn5 | contactin 5 | 7.629540386 | 0.033160908 |
| Matn1 | matrilin 1, cartilage matrix protein | 6.832374198 | 0.040018696 |
| Wnt7a | wingless-type MMTV integration site family, member 7A | 6.583543209 | 0.032273462 |
| Cd247 | CD247 antigen | 5.783789669 | 0.009023444 |
| Cntn2 | contactin 2 | 4.209958054 | 0.006978308 |
| Epha8 | Eph receptor A8 | 3.921338752 | 0.013368563 |
| Cntnap4 | contactin associated protein-like 4 | 3.861791156 | 0.000157213 |
| Lum | lumican | 3.689170664 | 4.71E-09 |
| Dcn | decorin | 3.221405678 | 0.000343832 |
| Dsg2 | desmoglein 2 | 3.056813863 | 0.002173833 |
| Fcna | ficolin A | 2.472419456 | 0.04380761 |
| Hhip | Hedgehog-interacting protein | 2.448442044 | 1.20E-15 |
| Acan | aggrecan | 2.425253241 | 0.007846407 |
| Grem1 | gremlin 1, DAN family BMP antagonist | 2.327684142 | 0.007236108 |
| Ptch2 | patched 2 | 2.249939294 | 3.26E-07 |
| Ncam2 | neural cell adhesion molecule 2 | 2.23694375 | 0.045071424 |
| Wnt2b | wingless-type MMTV integration site family, member 2B | 2.205026195 | 0.03750596 |
| Tmeff2 | transmembrane protein with EGF-like and two follistatin-like domains 2 | 2.139160406 | 0.001093468 |

B

| Gene | | FC v vs d | p-value |
|---|---|---|---|
| Npy | neuropeptide Y | 10.19250569 | 0.000948699 |
| Tmem179 | transmembrane protein 179 | 9.118002956 | 0.01718109 |
| Fgf23 | fibroblast growth factor 23 | 8.117354271 | 0.039466303 |
| Ndp | norrin cystine knot growth factor | 7.274089109 | 0.010072231 |
| Ntng2 | netrin G2 | 4.63821552 | 0.011953916 |
| Cdhr5 | cadherin-related family member 5 | 3.503448261 | 0.042958767 |
| Rarres1 | retinoic acid receptor responder (tazarotene induced) 1 | 3.34336286 | 0.026863539 |
| Rspo2 | R-spondin 2 | 3.302195912 | 0.029840352 |
| Cbln1 | cerebellin 1 precursor protein | 2.768737606 | 0.023099369 |
| Cbln2 | cerebellin 2 precursor protein | 2.667302524 | 2.73E-12 |
| Bmper | BMP-binding endothelial regulator | 2.461271596 | 4.53E-05 |
| Sct | secretin | 2.445677264 | 1.01E-07 |
| Sorcs1 | sortilin-related VPS10 domain containing receptor 1 | 2.430231212 | 0.001346064 |
| Ildr2 | immunoglobulin-like domain containing receptor 2 | 2.23190764 | 0.033426751 |
| Unc5c | Unc-5 netrin receptor C | 2.217657525 | 0.011686367 |
| Spon1 | spondin 1, (f-spondin) extracellular matrix protein | 2.142168556 | 1.17E-13 |
| Nell1 | NEL-like 1 | 2.090527715 | 0.000646551 |
| Gdf10 | growth differentiation factor 10 (bmp3b) | 2.022343931 | 0.000531037 |

**Table S1. Dorsal or ventral muscle genes identified by RNA-Sequencing,** Related to Figure 5

(A) Transmembrane and secreted proteins with known roles in cell signaling that are enriched >2-fold in dorsal myoblasts (one-way ANOVA, $p<0.05$).

(B) Transmembrane and secreted proteins with known roles in cell signaling that are enriched >2-fold in ventral myoblasts (one-way ANOVA, $p<0.05$).

A

| Gene | Forward primer | Reverse primer | Source |
|---|---|---|---|
| *vstm2b* | 5'-CGGAGCAAGGTAACAAATAAGG-3' | 5'-CTCAAGGGACTTGCTCAAGAGT-3' | Allen Brain Atlas |
| *pv* | 5'-TCTGCTCATCCAAGTTGCAG-3' | 5'-TCCTGAAGGACTCAACCCC-3' | Allen Brain Atlas |
| *trkc* (*ntrk3*) | 5'-TTGTCAAGTTCTATGGGGTGTG-3' | 5'-TAGTAGATTGGCTCCAACCAGG-3' | Allen Brain Atlas |

B

| Gene type | Gene | Catalog number |
|---|---|---|
| Muscle marker | *myod1* | 316081 |
| Mesenchymal marker | *prx1* | 485231 |
| Candidate: dorsal | *lum* | 480361 |
| Candidate: dorsal | *dcn* | 413281 |
| Candidate: ventral | *bmper* | 549561 |

**Table S2. Probes used in gene expression profiling,** Related to Figure 6

(A) Primers used to generate DIG- and FITC-conjugated probes for in situ hybridization. Sequences were obtained from Allen Brain Atlas (Lein et al., 2007).

(B) Probes purchased for RNAScope in situ hybridization.

**Figure S1.**

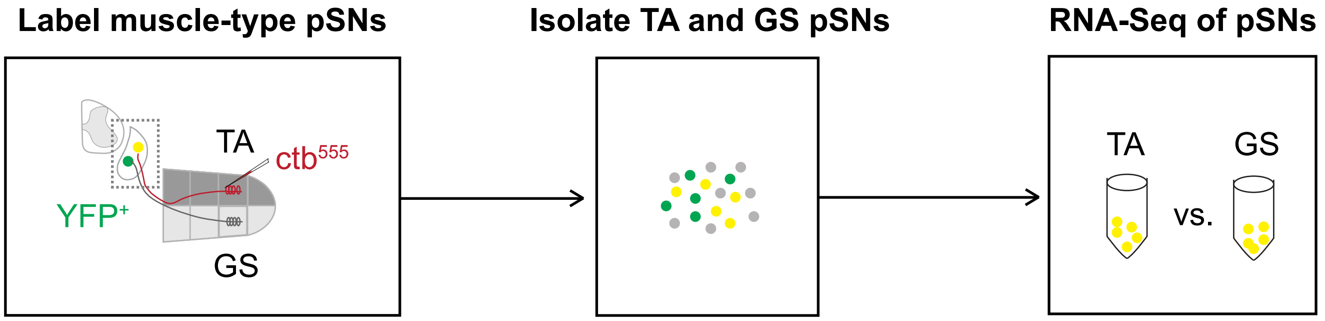


Inject TA or GS muscles of Pv::YFP mice with ctb at P0

Dissociate DRG at P1; aspirate individual YFP+, ctb+ neurons

1. Lyse pSNs
2. Prepare cDNA libraries
3. RNA-Sequencing
4. Compare gene expression

**Figure S2.**

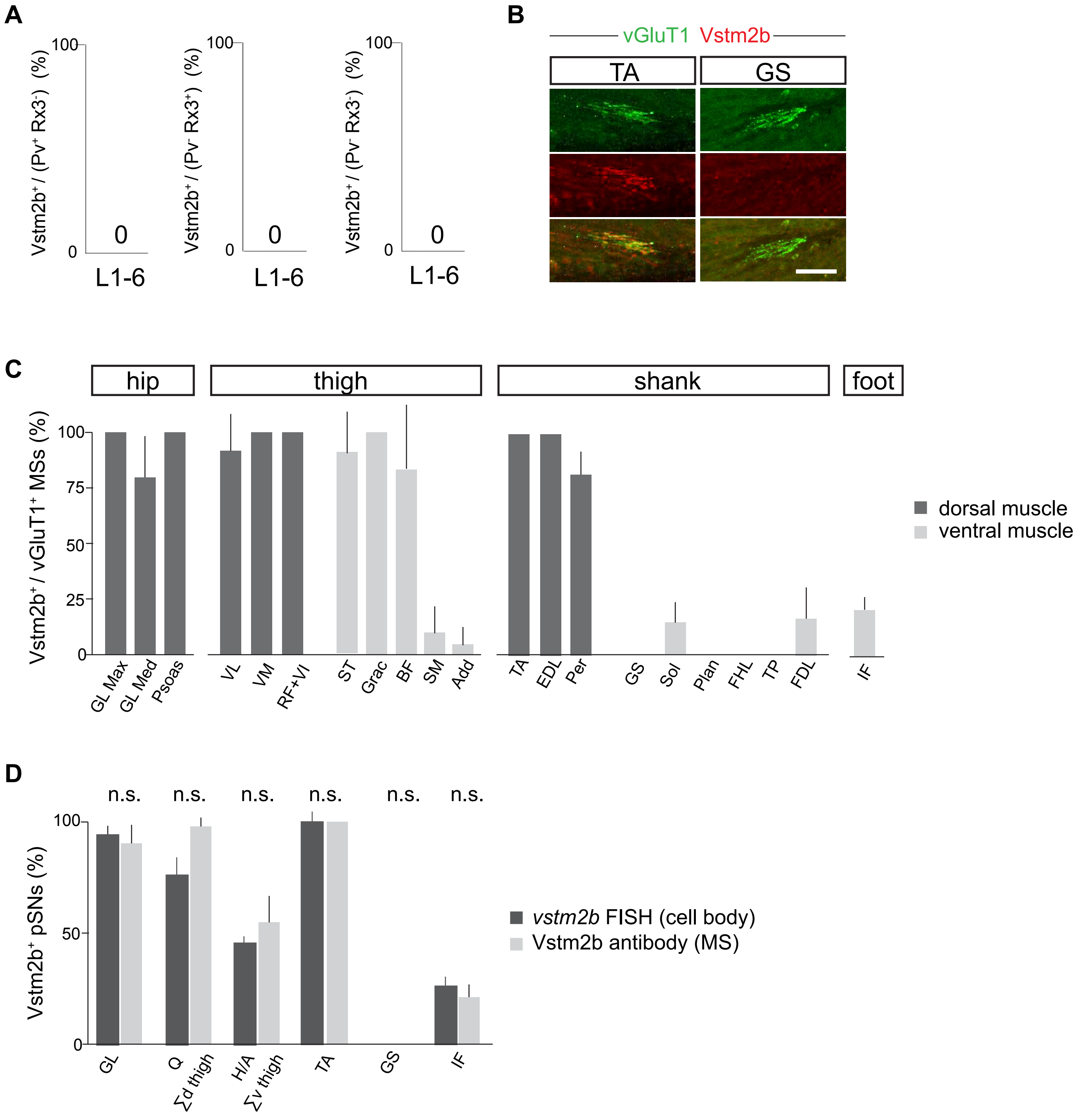
A
Vstm2b+ / (Pv+ Rx3-) (%)
Vstm2b+ / (Pv- Rx3+) (%)
Vstm2b+ / (Pv- Rx3-) (%)
100
0
L1-6
B
vGluT1 Vstm2b
TA
GS
C
hip
thigh
shank
foot
Vstm2b+ / vGluT1+ MSs (%)
100
75
50
25
0
GL Max
GL Med
Psoas
VL
VM
RF+VI
ST
Grac
BF
SM
Add
TA
EDL
Per
GS
Sol
Plan
FHL
TP
FDL
IF
dorsal muscle
ventral muscle
D
n.s.
Vstm2b+ pSNs (%)
100
50
0
GL
Q
Σd thigh
H/A
Σv thigh
TA
GS
IF
vstm2b FISH (cell body)
Vstm2b antibody (MS)

**Figure S3.**

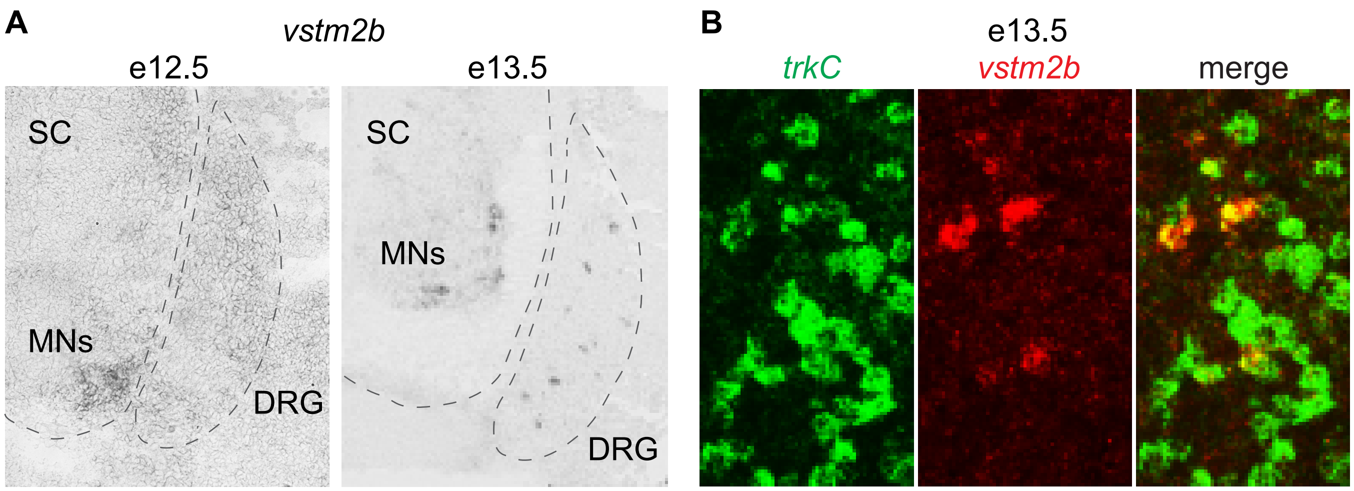
A
vstm2b
e12.5
e13.5
SC
MNs
DRG
SC
MNs
DRG
B
e13.5
trkC
vstm2b
merge

**Figure S4.**

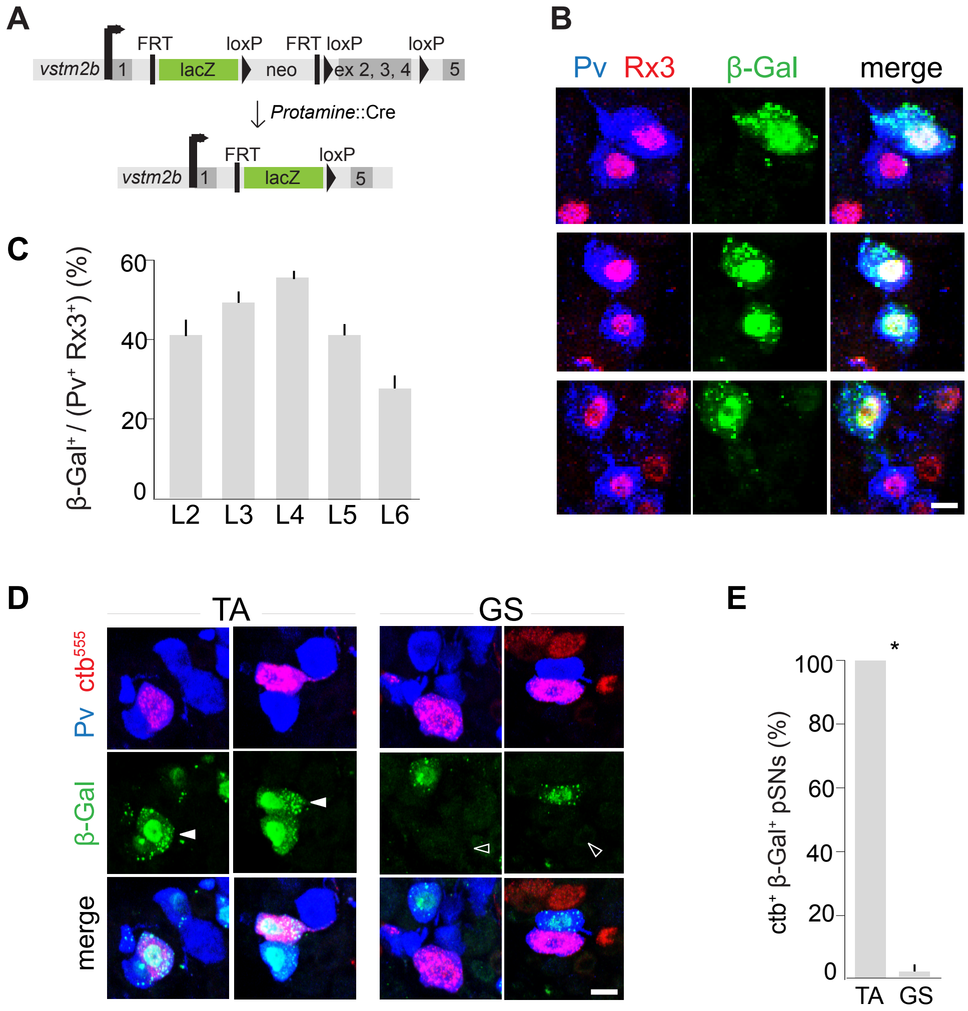
A
vstm2b
1
FRT
lacZ
loxP
neo
FRT loxP
ex 2, 3, 4
loxP
5
Protamine::Cre
vstm2b
1
FRT
lacZ
loxP
5
B
Pv Rx3
β-Gal
merge
C
β-Gal+ / (Pv+ Rx3+) (%)
60
40
20
0
L2 L3 L4 L5 L6
D
TA
GS
Pv ctb555
β-Gal
merge
E
ctb+ β-Gal+ pSNs (%)
100
80
60
40
20
0
*
TA GS

**Figure S5.**

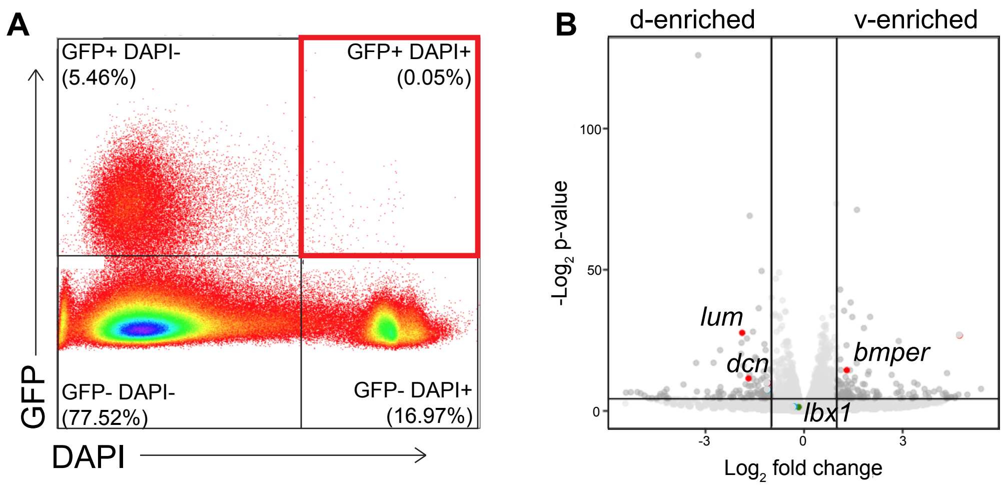
A
GFP+ DAPI-
(5.46%)
GFP+ DAPI+
(0.05%)
GFP- DAPI-
(77.52%)
GFP- DAPI+
(16.97%)
GFP
DAPI
B
d-enriched
v-enriched
100
50
0
-Log2 p-value
lum
dcn
bmper
lbx1
-3
0
3
Log2 fold change

**Figure S6.**

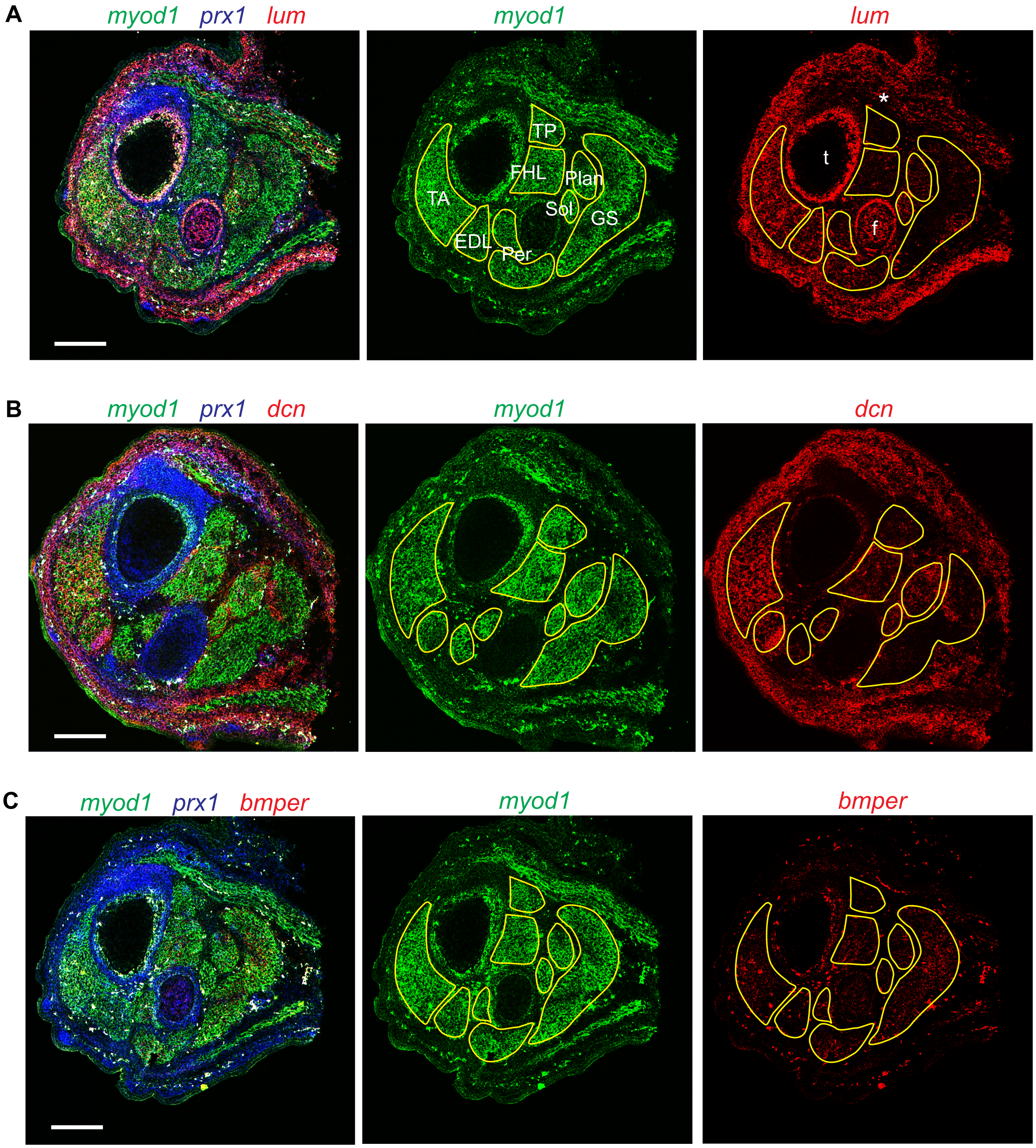

A
myod1 prx1 lum
myod1
TP
FHL
Plan
TA
Sol
GS
EDL
Per
lum
*
t
f
B
myod1 prx1 dcn
myod1
dcn
C
myod1 prx1 bmper
myod1
bmper
